# Polygenic and clinical risk scores and their impact on age at onset of cardiometabolic diseases and common cancers

**DOI:** 10.1101/727057

**Authors:** Nina J. Mars, Jukka T. Koskela, Pietari Ripatti, Tuomo T.J. Kiiskinen, Aki S. Havulinna, Joni V. Lindbohm, Ari Ahola-Olli, Mitja Kurki, Juha Karjalainen, Priit Palta, FinnGen, Benjamin M. Neale, Mark Daly, Veikko Salomaa, Aarno Palotie, Elisabeth Widén, Samuli Ripatti

## Abstract

**Background:** Polygenic risk scores (PRS) have shown promise in predicting susceptibility to common diseases. However, the extent to which PRS and clinical risk factors act jointly and identify high-risk individuals for early onset of disease is unknown.

**Methods:** We used large-scale biobank data (the FinnGen study; n=135,300), with up to 46 years of prospective follow-up, and the FINRISK study with standardized clinical risk factor measurements to build genome-wide PRSs with >6M variants for coronary heart disease (CHD), type 2 diabetes (T2D), atrial fibrillation (AF), and breast and prostate cancer. We evaluated their associations with first disease events, age at disease onset, and impact together with routinely used clinical risk scores for predicting future disease.

**Results:** Compared to the 20-80^th^ percentiles, a PRS in the top 2.5% translated into hazard ratios (HRs) for incident disease ranging from 2.03 to 4.28 (p-values 1.96×10^−59^ to <1.00×10^−100^) and the bottom 2.5% into HRs ranging from 0.20 to 0.61. The estimated difference in age at disease onset between top and bottom 2.5% of PRSs was 6 to 13 years. Among early-onset cases, 21.3-32.9% had a PRS in the highest decile and in CHD and AF.

**Conclusions:** The properties of PRS were similar in all five diseases. PRS identified a considerable proportion early-onset cases, and for all ages the performance of PRS was comparable to established clinical risk scores. These findings warrant further clinical studies on application of polygenic risk information for stratified screening or for guiding lifestyle and preventive medical interventions.

## INTRODUCTION

Common chronic diseases present a huge burden to societies, with an estimated one billion prevalent cases diagnosed with cardiovascular diseases, diabetes, or neoplasms worldwide.^1^Consequently, the development of strategies to prevent these diseases is critically important. To facilitate prevention, a clear understanding of individual risk is essential to determine whether an individual warrants an intervention as well as to gauge the impact of different interventions. These risk models typically incorporate clinical and laboratory-based risk factors, and can identify individuals at high risk suitable for selective prevention strategies, such as prescribing cholesterol-lowering medications for reducing coronary heart disease (CHD) risk.^2^ Although clinical risk scores enable the identification of individuals who may benefit from preventive interventions, these models are imperfect and fail to identify a noticeable proportion of those who develop the diseases, particularly among young individuals.^2,3^ Furthermore, in many cancers such as breast and prostate cancer, no widely accepted clinical risk assessment strategy exists.

Large-scale genetic screens comparing disease cases with controls have identified thousands of genetic loci associated with risk of complex diseases,^4^suggesting that genomic information has become a promising candidate for improving clinical risk assessment.^5^While, individually, the associated loci typically modify the disease risks only marginally, for many diseases the cumulative impact of risk across the genome is considerable.^6^Polygenic risk scores (PRS) measure these cumulative genetic effects^7^and have recently been shown to correlate with cases status in many complex diseases including CHD, T2D, and breast cancer.^8-10^ However, limited information exists as to both the performance of PRSs in a prospective setting and their value when integrated with the established clinical risk factors and laboratory-based biomarkers.

We set to test the utility of PRSs derived from large-scale genomic information for predicting first disease events in five diseases: CHD, T2D, atrial fibrillation or flutter (AF), and breast and prostate cancer. Specifically, we tested three hypotheses: 1) are the polygenic risk scores associated with first disease events over a long follow-up, 2) can polygenic risk scores help identify early-onset cases, and 3) what is the relative impact of clinical risk scores and the polygenic risk scores? We tested these hypotheses within the FinnGen study cohort comprising of 135,300 individuals with genome-wide genotyping and up to 46 years of follow-up.

## METHODS

### Individuals

The data comprised of 135,300 Finnish individuals from FinnGen Preparatory Phase Data Freeze 3, which includes prospective epidemiological and disease-based cohorts, and hospital biobank samples (Supplementary Table S1). The data, representing roughly 3% of Finnish adult population, were linked by the unique national personal identification numbers to national hospital discharge (available from 1968), death (1969-), cancer (1953-), and medication reimbursement (1995-) registries.

A subset of FinnGen, the population-based FINRISK study with 21,813 individuals was selected for analyzing the PRSs together with clinical risk factors. The FINRISK surveys, performed in 1992, 1997, 2002, and 2007 comprised random samples of adults within five geographical areas in Finland. The baseline data covered self-reported information assessed by questionnaires, anthropometric measurements, and blood samples. Additional details on the study protocol have been previously described.^11^ Information on the genotyping and imputation are provided in the Supplemental Appendix. The study was approved by the Coordinating Ethics Committee of the Hospital District of Helsinki and Uusimaa (HUS/990/2017). The FINRISK analyses were conducted using the THL biobank permission for project BB2015_55.1. All participants gave written informed consent.

### Disease endpoints

Using the national registries, we studied the incidence of five diseases: CHD, T2D, AF, breast cancer, and prostate cancer (diagnoses based on International Classification of Diseases, ICD-8, ICD-9, and ICD-10 in Supplementary Table S2). Follow-up ended at first-ever diagnosis of the disease of interest, death, or at the end of follow-up on December 31, 2015, whichever came first.

### Clinical risk factors

Clinical risk factors were evaluated for CHD, T2D, and AF. The 10-year risk of hard atherosclerotic cardiovascular disease (ASCVD) was evaluated with the pooled cohort equations (PCE) according to guidelines,^12,13^ comprising age, sex, self-reported ancestry, total cholesterol (TC), high-density lipoprotein (HDL), systolic blood pressure (SBP), blood pressure-lowering medication, diabetes, and smoking status. The 10-year risk for ASCVD was categorized as intermediate to high risk (≥7.5% which often leads to consideration of preventive medication) or low to borderline risk (<7.5%).^13^

For T2D, individuals who had impaired fasting glucose (≥5.6 mmol/L) or who met the American Diabetes Association (ADA) criteria for testing for diabetes or prediabetes in asymptomatic adults^14^ were defined as having high clinical risk for T2D. Family history was the self-reported parental history for any diabetes and early myocardial infarction (MI). Following many previous studies, the definition for early-onset cases was age 55 for CHD, 45 for T2D, 60 for AF, 40 for breast cancer, and 55 for prostate cancer.

For AF, the clinical assessment of the 5-year absolute risk for individuals above age 45 was carried out with the CHARGE-AF score, comprising age, height, weight, systolic and diastolic blood pressure, smoking status, blood pressure-lowering treatment, prevalent diabetes, heart failure, and history of myocardial infarction.^15^ CHARGE-AF was revised and the risk was categorized as ≤5% or >5%.^15^ Additional information on the clinical risk factors are provided in the Supplemental Appendix.

### Polygenic risk scores

The summary association statistics came from recent genome-wide association studies (GWAS) (Supplementary Table S3, Supplementary Figure 1).^16-20^ LDpred was used to account for linkage disequilibrium among loci,^21^with whole-genome sequencing data on 2,690 Finns serving as the LD reference panel. After performing quality control, the final scores were generated with PLINK2^24^by calculating the weighted sum of risk allele dosages for each single nucleotide polymorphism (SNP). The final PRSs comprised 6,412,950 variants for CHD PRS, 6,437,380 for T2D PRS, 6,171,733 for AF PRS, 6,390,808 for breast cancer PRS, and 6,606,785 for prostate cancer PRS (candidate LDpred scores with respect to the tuning parameter in Supplementary Table S4).

### Statistical analysis

The PRS distributions were divided into bins of <2.5, 2.5-20, 20-80, 80-97.5, >97.5, using 20-80 as the reference, to display the risk in the tails with respect to a large group of individuals defined to have average risk. When comparing clinical and polygenic risk, we defined elevated PRS as a polygenic riskscore above the 90^th^percentile. Cox proportional hazards model was used to estimate survival curves and hazard ratios (HRs) and 95% confidence intervals (CI). Schoenfeld residuals and log-log inspection showed that proportional assumption criteria applied in our models. Unless otherwise stated, we used age as the time scale, stratified regression models by sex and adjusted for FINRISK survey collection year, genotyping array, and the first ten principal components of ancestry. Breast cancer was studied only in women and prostate cancer only in men. The relative contributions of PRS and clinical scores were estimated using logistic regression and discrimination was evaluated by the area under the receiver operating characteristic curve (AUC), and for the survival models by C-index.

In FINRISK analyses, the association of PRSs was tested for incident disease cases only. Participants with prevalent diabetes were excluded from all prospective CHD analyses performed in FINRISK. The FINRISK had 1805 individuals overlapping with the AF GWAS, and we excluded these individuals from the AF analyses. Age at disease onset and the differences between PRS categories were estimated with restricted mean survival time (RMST).^22^ For statistical analyses, we used R 3.5.2, and Stata 14.2 (College Station, TX, USA).

## RESULTS

### Polygenic risk score associations

We first derived PRSs for five diseases, CHD, T2D, AF, breast cancer, and prostate cancer by weighting the individual SNPs by their effect sizes from published GWASs and by accounting for linkage disequilibrium between markers. We tested the association between these newly derived PRSs and disease events within the independent FinnGen study cohort (n = 135,300), which comprised 20,179 individuals with CHD, 17,519 with T2D, 12,809 with AF, 3,904 with breast cancer, and 3,263 with prostate cancer. FinnGen was comprised of 56.3% women, with mean age 59.2 (standard deviation, SD 16.6) at the end of follow-up.

For all five diseases, a higher PRS was strongly associated with a higher incidence rate (Figure 1, Table 1). The HR per SD increment (all p-values <1.00×10^−100^) was 1.31 for CHD (95% CI 1.29-1.33), for T2D 1.74 (1.72-1.77), for AF 1.62 (1.59-1.65), for breast cancer 1.67 (1.62-1.72), and for prostate cancer 1.86 (1.80-1.93). Compared to individuals with average PRS (20-80^th^ percentile of the PRS distribution), being in the top 2.5% of the distribution translated into HRs ranging from 2.03 to 4.28 (p-values 1.96×10^−59^to <1.00×10^−100^; Table 1). Similarly, in the lowest 2.5% of the PRS, the HRs when comparing to the average PRS were 0.20-0.61 (p-values 4.35×10^−10^ to 7.11×10^−64^).

**Figure 1.**
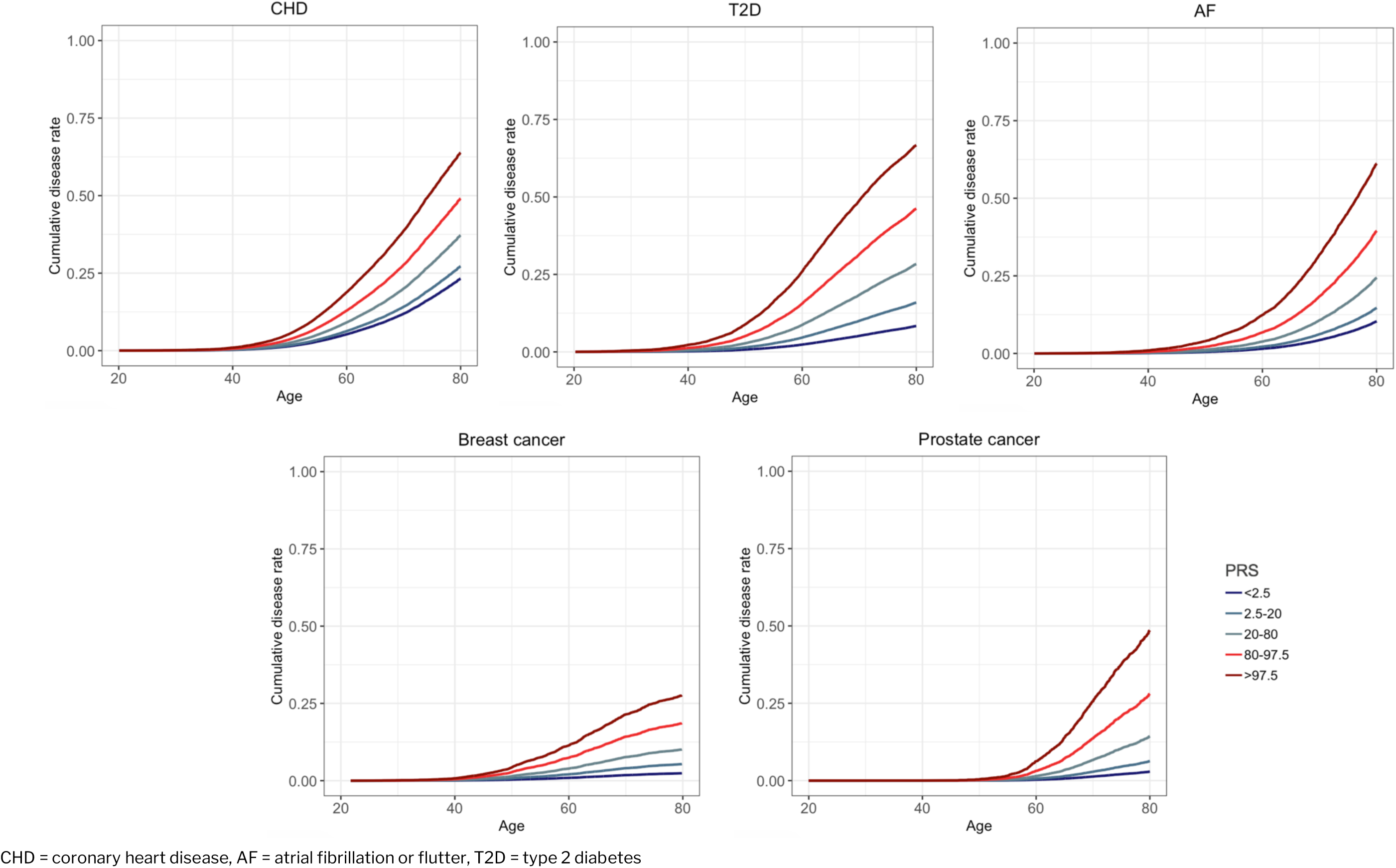
Adjusted survival curves in FinnGen, showing the cumulative risk of disease by polygenic risk score (PRS) categories (p-value for trend <1.00×10^−100^ for each).

**Table 1.** Hazard ratios (HR) and 95% confidence intervals (CI) for polygenic risk score (PRS) bins in FinnGen.

|  | HR (95% CI) | p | Cases / Controls |
| --- | --- | --- | --- |
| **CHD PRS** |  |  |  |
| <2.5 | 0.61 (0.55-0.69) | $3.04 \times 10^{-18}$ | 329 / 3,054 |
| 2.5-20 | 0.72 (0.69-0.75) | $2.41 \times 10^{-53}$ | 2,580 / 21,097 |
| 20-80 | 1 (reference) | - | 11,832 / 69,348 |
| 80-97.5 | 1.38 (1.34-1.43) | $1.30 \times 10^{-76}$ | 4,532 / 19,145 |
| >97.5 | 2.03 (1.90-2.18) | $2.87 \times 10^{-93}$ | 906 / 2,477 |
| **T2D PRS** |  |  |  |
| <2.5 | 0.22 (0.19-0.27) | $7.11 \times 10^{-64}$ | 131 / 3,195 |
| 2.5-20 | 0.48 (0.46-0.51) | $<1.00 \times 10^{-100}$ | 1,653 / 21,548 |
| 20-80 | 1 (reference) | - | 9,869 / 68,998 |
| 80-97.5 | 1.92 (1.85-1.99) | $<1.00 \times 10^{-100}$ | 4,776 / 17,970 |
| >97.5 | 3.45 (3.24-3.67) | $<1.00 \times 10^{-100}$ | 1,090 / 2,072 |
| **AF PRS** |  |  |  |
| <2.5 | 0.39 (0.33-0.46) | $1.34 \times 10^{-27}$ | 136 / 3,247 |
| 2.5-20 | 0.56 (0.53-0.59) | $3.92 \times 10^{-81}$ | 1,281 / 22,396 |
| 20-80 | 1 (reference) | - | 7,159 / 74,021 |
| 80-97.5 | 1.81 (1.73-1.88) | $<1.00 \times 10^{-100}$ | 3,417 / 20,260 |
| >97.5 | 3.50 (3.26-3.77) | $<1.00 \times 10^{-100}$ | 816 / 2,567 |
| **Breast cancer PRS** |  |  |  |
| <2.5 | 0.23 (0.15-0.35) | $1.27 \times 10^{-11}$ | 21 / 1,885 |
| 2.5-20 | 0.52 (0.46-0.59) | $1.45 \times 10^{-27}$ | 320 / 13,020 |
| 20-80 | 1 (reference) | - | 2,126 / 43,606 |
| 80-97.5 | 1.90 (1.77-2.04) | $1.33 \times 10^{-69}$ | 1,179 / 12,160 |
| >97.5 | 2.93 (2.58-3.34) | $1.96 \times 10^{-59}$ | 258 / 1,648 |
| **Prostate cancer PRS** |  |  |  |
| <2.5 | 0.20 (0.12-0.33) | $4.35 \times 10^{-10}$ | 15 / 1,462 |
| 2.5-20 | 0.43 (0.37-0.49) | $3.63 \times 10^{-32}$ | 215 / 10,123 |
| 20-80 | 1 (reference) | - | 1,725 / 33,716 |
| 80-97.5 | 2.12 (1.96-2.29) | $1.08 \times 10^{-79}$ | 1,020 / 9,317 |
| >97.5 | 4.28 (3.77-4.85) | $<1.00 \times 10^{-100}$ | 288 / 1,188 |

CHD = coronary heart disease, AF = atrial fibrillation or flutter, T2D = type 2 diabetes

In addition, we built estrogen receptor-negative (ER-negative) and estrogen receptor-positive (ER-positive) breast cancer PRSs (Supplementary Figure 2). With any breast cancer as the outcome, for ER-negative PRS the HR for average PRS vs top 2.5% of the PRS distribution was 1.77 (95% CI 1.51-2.07, p = 9.00×10^−13^) and for ER-positive PRS, 2.75 (95% CI 2.41-3.13, p = 1.26×10^−50^).

The higher the PRS, the earlier was the disease onset for all five diseases (Figure 2, sex-specific results in Supplementary Figure 3). Compared to individuals with average PRS, those in the top 2.5% of the distribution had a disease onset 4 to 9 years earlier. The largest difference in age at onset between the top and bottom 2.5%, 13.4 years, was seen for T2D.

**Figure 2.**
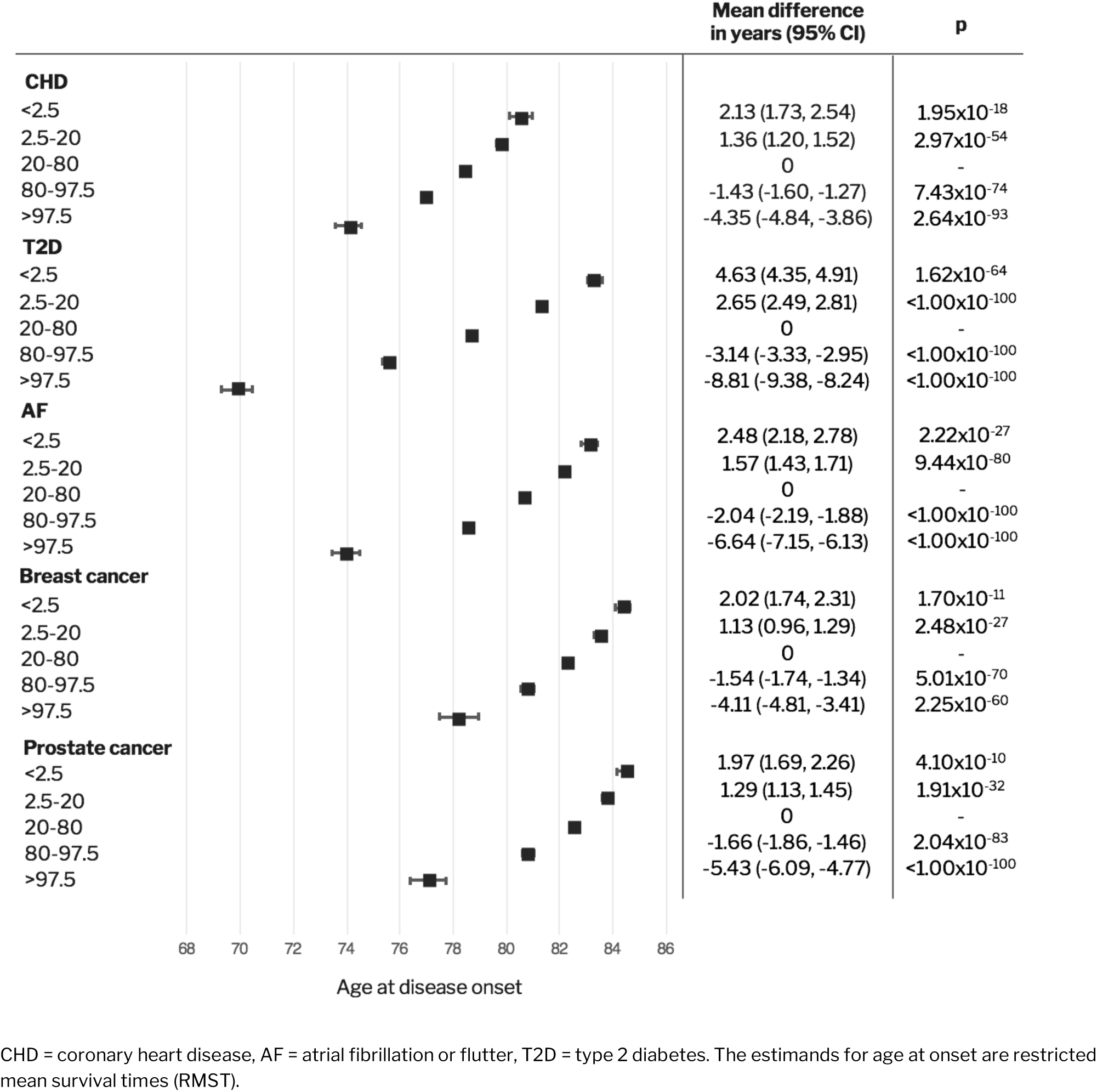
Difference in ages at disease onset across polygenic risk score categories in FinnGen.

### Relative contribution of polygenic and clinical risk for CHD, T2D, and AF

In CHD and T2D, clinical risk factors are routinely used for risk evaluation. Therefore, we assessed the relative and combined effect of PRSs and clinical risk calculators, using the FINRISK study (n = 21,813, mean age at baseline 48.0, 52.7% women) with major cardiometabolic risk factors measured (Supplementary Table S5). Overall, the correlation between polygenic and clinical risk scores was weak, and family history of CHD or T2D had only a minor effect on the association between polygenic risk and disease (Supplementary Figure 4; Supplementary Table S6).

To guide decision-making, prevention guidelines report thresholds for clinical risk calculators. These thresholds were used for identifying individuals at clinical high risk, and to assess the extent to which PRS and clinical risk assessment individually and jointly identify high-risk individuals. PRS helped identify individuals at very high risk: in those with both high clinical risk and high PRS, the risk was elevated compared to those with only high clinical risk or only high PRS (Figure 3). Combining both scores provided only a minor increment to the PRS accuracy (Supplementary Figure 5).

**Figure 3.**
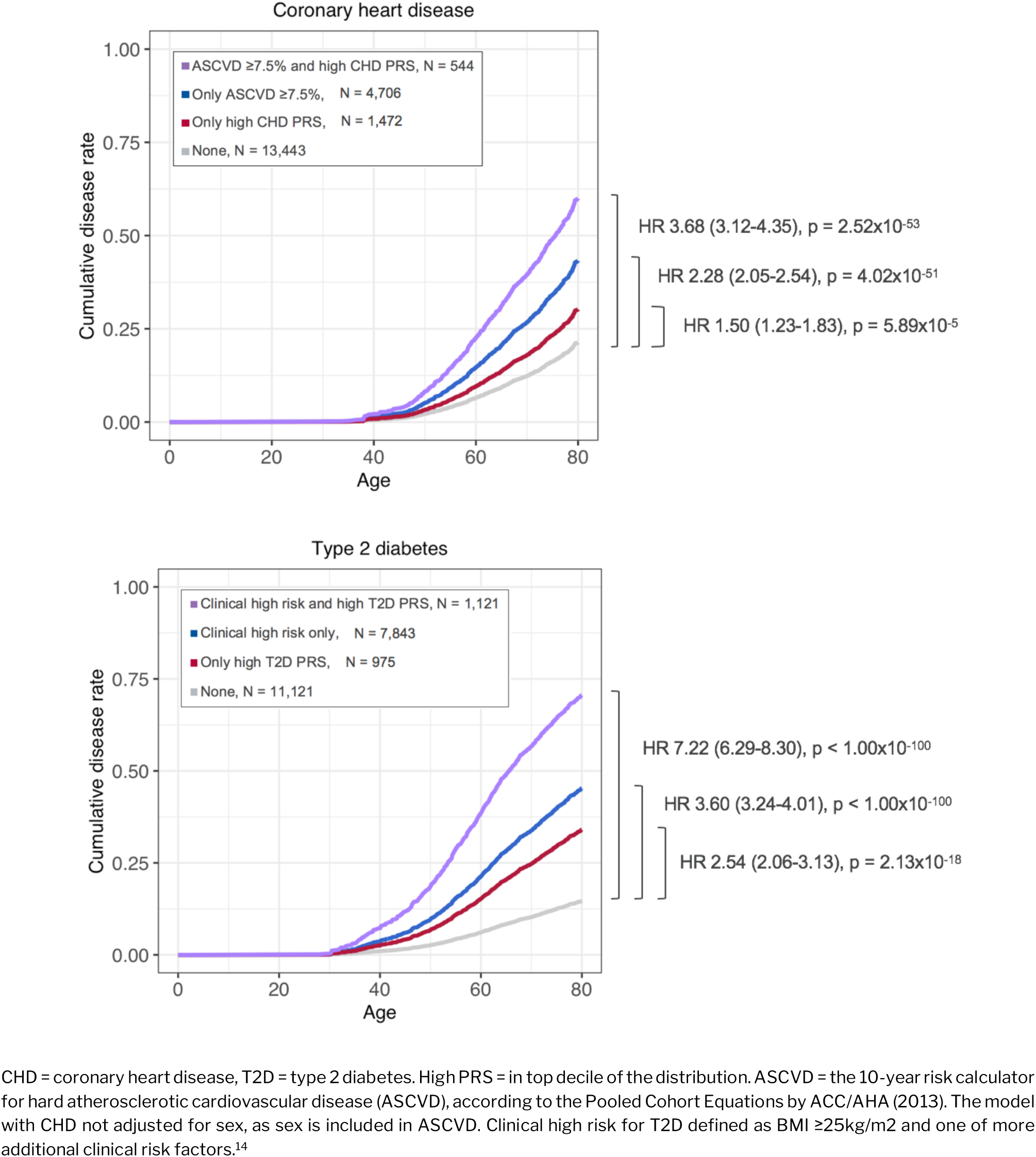
Adjusted survival curves with respective hazard ratios (95% confidence interval) in FINRISK for high clinical risk, high PRS, and both.

After dividing the individuals into early- and late-onset cases, we compared how often either the PRS or clinical risk was elevated among cases. In all five diseases, around one-quarter of the early-onset cases also had an elevated PRS, whereas around one-fifth of the late-onset cases had an elevated PRS (Figure 4). Compared to early-onset cases, a larger proportion of late-onset cases had high clinical risk. In early-onset CHD, the proportion of either elevated PRS or ASCVD ≥7.5% was 39.7% and 6.7% had both risk scores elevated. In early-onset T2D, high clinical risk was over twice more common than an elevated PRS, with 73.2% having one risk score elevated and 14.4% having both scores elevated. In early-onset AF, 25.0% had an elevated PRS, whereas hardly anyone had CHARGE-AF >5%.

**Figure 4.**
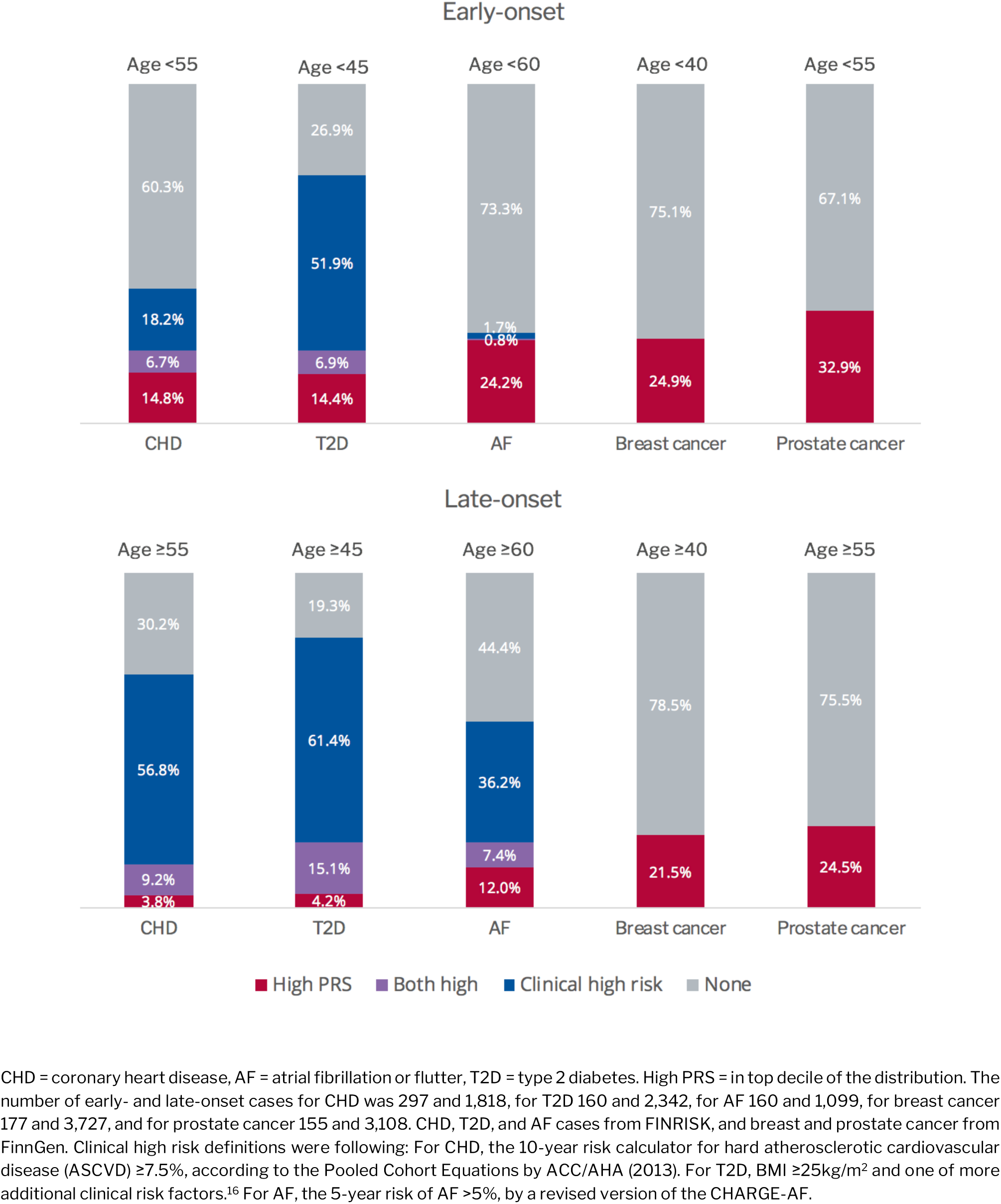
The proportion early- and late-onset cases with high clinical risk, high polygenic risk, or neither.

### Effect of polygenic risk on breast and prostate cancer prevalence

Breast cancer screening starts from age 50 in Finland, when its cumulative incidence reaches 2.0%. Using the population-based FINRISK study, we estimated the age when the cumulative incidence reached 2.0% in the different PRS groups: 44.5 years in PRS category >97.5%, 45.4 years in 80-97.5%, 50.0 years in 20-80%, and 58.5 years in PRS <20%. Similarly, a 2% cumulative incidence for prostate cancer in men was reached at age 62. When estimated separately for the PRS groups, a 2% cumulative incidence was reached at age 55.6 in PRS category >97.5%, at 59.4 in 80-97.5%, at 62.4 in 20-80%, and at 69.9 in <20%.

## DISCUSSION

For all five common diseases, we detected similar patterns for the impact of polygenic risk. Over the long follow-up, high polygenic risk correlated with a substantial elevation in disease risk, replicating the results from earlier cross-sectional investigations applying highly polygenic scores,^8^ now in a population-based setting with population-level disease prevalence and decades-long follow-up. This risk elevation also translated into large shifts in ages at disease onset. Compared to individuals with an average PRS, those in the top 2.5% of the PRS distribution had a disease onset 4 to 9 years earlier. Of early-onset cases, approximately one-quarter had an elevated PRS.

The relative contribution of polygenic and clinical risk was assessed for CHD, T2D, and AF. The PRS complemented routinely used clinical risk calculators and identified high-risk individuals missed by established clinical risk calculators. As most cardiovascular risk calculators have been trained with data on middle-aged individuals, their ability to identify individuals at risk for early-onset CHD is limited.^2,12^ In early-onset CHD and AF, the PRS identified more high-risk individuals than did the routinely used clinical risk calculators, but in all the diseases, the PRS influenced disease risk also in late-onset cases, and improved identification of those at very high disease risk. Hence, polygenic and clinical risk scores complement each other and provide most benefit for prediction when utilized together. As early disease onset may also implicate more aggressive forms of disease,^23,24^ preventing or postponing the disease onset in individuals with high polygenic risk could translate into considerable health benefits. How to integrate PRS into clinical practice and how to communicate this information to patients should carefully be assessed for each disease.

In many countries including Finland, breast cancer screening is initiated at around age 50 by which approximately 2% of women have been diagnosed with breast cancer.^25,26^ In our data, this 2% prevalence was reached at age 50 in those with an average PRS, with the difference between the tails of the PRS distribution amounting to 14 years. Our results suggest that PRS could provide new tools for risk stratification even in situations where only few robust clinical risk factors exist, as in the case of the common cancers assessed in this study.

As the data comprised of individuals of European ancestry, PRSs need to be tested also in non-European samples.^27^ It is of utmost importance to conduct GWASs in non-Europeans to provide input for polygenic risk scores in populations of non-European origin.^28^ Although our analyses were performed in Finns, our results are in line with earlier reports from other samples of European origin.^8,9,29^ A fraction of individuals in FinnGen were ascertained through hospital biobanks or disease-based cohorts, which may lead to overestimation of risks. However, the effects of the PRSs in FinnGen were highly similar to those in the population-based FINRISK (Supplementary Tables S7-S8, Supplementary Figure 6). Reasons for selecting this range of diseases were twofold. First, we aimed to explore patterns for polygenic risk across a variety of common diseases. Second, each have large existing GWAS with openly accessible summary-level data for generating these highly polygenic scores. Similar patterns are expected to be present in other common and heritable diseases.^30^

In conclusion, when predicting first disease events, polygenic risk scores identified individuals missed by established clinical risk prediction models, particularly those at high risk for early-onset disease. The opportunities to apply polygenic risk information for stratified screening or for guiding lifestyle and medical interventions in the clinical setting remain to be defined in further studies.

## Acknowledgements

We would like to thank Sari Kivikko, Huei-Yi Shen, and Ulla Tuomainen for management assistance. For the Finnish Institute of Health and Welfare (THL) driven FinnGen preparatory project (here called FinnGen), all patients and control subjects had provided informed consent for biobank research, based on the Finnish Biobank Act. Alternatively, older cohorts were based on study-specific consents and later transferred to the THL Biobank after approval by Valvira, the National Supervisory Authority for Welfare and Health. Recruitment protocols followed the biobank protocols approved by Valvira. The Ethical Review Board of the Hospital District of Helsinki and Uusimaa approved the FinnGen study protocol Nr HUS/990/2017. The FinnGen preparatory project is approved by THL, approval numbers THL/2031/6.02.00/2017, amendments THL/341/6.02.00/2018, THL/2222/6.02.00/2018 and THL/283/6.02.00/2019. The FINRISK analyses were conducted using the THL biobank permission for project BB2015_55.1. The FINRISK data used for the research were obtained from THL Biobank. We thank all study participants for their generous participation in FINRISK and FinnGen. The content is solely the responsibility of the authors and does not necessarily represent the official views of the National Institutes of Health.

## REFERENCES

1. Global Burden of Disease. Global, regional, and national incidence, prevalence, and years lived with disability for 354 diseases and injuries for 195 countries and territories, 1990-2017: a systematic analysis for the Global Burden of Disease Study 2017. Lancet 2018;392:1789–858.

2. Piepoli MF, Hoes AW, Agewall S, et al. 2016 European Guidelines on cardiovascular disease prevention in clinical practice: The Sixth Joint Task Force of the European Society of Cardiology and Other Societies on Cardiovascular Disease Prevention in Clinical Practice (constituted by representatives of 10 societies and by invited experts)Developed with the special contribution of the European Association for Cardiovascular Prevention & Rehabilitation (EACPR). Eur Heart J 2016;37:2315–81.

3. Lindbohm JV, Sipila PN, Mars NJ, et al. 5-year versus risk-category-specific screening intervals for cardiovascular disease prevention: a cohort study. Lancet Public Health 2019;4:e189–e99.

4. Visscher PM, Wray NR, Zhang Q, et al. 10 Years of GWAS Discovery: Biology, Function, and Translation. Am J Hum Genet 2017;101:5–22.

5. Torkamani A, Wineinger NE, Topol EJ. The personal and clinical utility of polygenic risk scores. Nat Rev Genet 2018.

6. Timpson NJ, Greenwood CMT, Soranzo N, Lawson DJ, Richards JB. Genetic architecture: the shape of the genetic contribution to human traits and disease. Nat Rev Genet 2018;19:110–24.

7. Chatterjee N, Shi J, Garcia-Closas M. Developing and evaluating polygenic risk prediction models for stratified disease prevention. Nat Rev Genet 2016;17:392–406.

8. Khera AV, Chaffin M, Aragam KG, et al. Genome-wide polygenic scores for common diseases identify individuals with risk equivalent to monogenic mutations. Nat Genet 2018;50:1219–24.

9. Mavaddat N, Michailidou K, Dennis J, et al. Polygenic Risk Scores for Prediction of Breast Cancer and Breast Cancer Subtypes. Am J Hum Genet 2019;104:21–34.

10. Abraham G, Havulinna AS, Bhalala OG, et al. Genomic prediction of coronary heart disease. Eur Heart J 2016;37:3267–78.

11. Borodulin K, Tolonen H, Jousilahti P, et al. Cohort Profile: The National FINRISK Study. Int J Epidemiol 2017.

12. Goff DC, Jr., Lloyd-Jones DM, Bennett G, et al. 2013 ACC/AHA guideline on the assessment of cardiovascular risk: a report of the American College of Cardiology/American Heart Association Task Force on Practice Guidelines. Circulation 2014;129:S49–73.

13. Grundy SM, Stone NJ, Bailey AL, et al. 2018 AHA/ACC/AACVPR/AAPA/ABC/ACPM/ADA/AGS/APhA/ASPC/NLA/PCNA Guideline on the Management of Blood Cholesterol: A Report of the American College of Cardiology/American Heart Association Task Force on Clinical Practice Guidelines. J Am Coll Cardiol 2018.

14. American Diabetes Association. 2. Classification and Diagnosis of Diabetes: Standards of Medical Care in Diabetes-2019. Diabetes Care 2019;42:S13–S28.

15. Alonso A, Krijthe BP, Aspelund T, et al. Simple risk model predicts incidence of atrial fibrillation in a racially and geographically diverse population: the CHARGE-AF consortium. J Am Heart Assoc 2013;2:e000102.

16. Zhou W, Nielsen JB, Fritsche LG, et al. Efficiently controlling for case-control imbalance and sample relatedness in large-scale genetic association studies. Nat Genet 2018;50:1335–41.

17. Mahajan A, Taliun D, Thurner M, et al. Fine-mapping type 2 diabetes loci to single-variant resolution using high-density imputation and islet-specific epigenome maps. Nat Genet 2018;50:1505–13.

18. Schumacher FR, Al Olama AA, Berndt SI, et al. Association analyses of more than 140,000 men identify 63 new prostate cancer susceptibility loci. Nat Genet 2018;50:928–36.

19. Michailidou K, Lindstrom S, Dennis J, et al. Association analysis identifies 65 new breast cancer risk loci. Nature 2017;551:92–4.

20. Nielsen JB, Thorolfsdottir RB, Fritsche LG, et al. Biobank-driven genomic discovery yields new insight into atrial fibrillation biology. Nat Genet 2018;50:1234–9.

21. Vilhjalmsson BJ, Yang J, Finucane HK, et al. Modeling Linkage Disequilibrium Increases Accuracy of Polygenic Risk Scores. Am J Hum Genet 2015;97:576–92.

22. Royston P, Parmar MK. Restricted mean survival time: an alternative to the hazard ratio for the design and analysis of randomized trials with a time-to-event outcome. BMC Med Res Methodol 2013;13:152.

23. Salinas CA, Tsodikov A, Ishak-Howard M, Cooney KA. Prostate cancer in young men: an important clinical entity. Nat Rev Urol 2014;11:317–23.

24. Sharifi M, Higginson E, Bos S, et al. Greater preclinical atherosclerosis in treated monogenic familial hypercholesterolemia vs. polygenic hypercholesterolemia. Atherosclerosis 2017;263:405–11.

25. Average Number of New Cases Per Year and Age-Specific Incidence Rates per 100,000 Females, UK. https://www.cancerresearchuk.org/health-professional/cancer-statistics/statistics-by-cancer-type/breast-cancer Accessed 4 April, 2019.

26. NORDCAN: Cancer Incidence, Mortality, Prevalence and Survival in the Nordic Countries, Version 8.2 (26.03.2019). http://www-dep.iarc.fr/NORDCAN/ Accessed 4 April, 2019.

27. Martin AR, Gignoux CR, Walters RK, et al. Human Demographic History Impacts Genetic Risk Prediction across Diverse Populations. Am J Hum Genet 2017;100:635–49.

28. Martin AR, Kanai M, Kamatani Y, Okada Y, Neale BM, Daly MJ. Clinical use of current polygenic risk scores may exacerbate health disparities. Nat Genet 2019;51:584–91.

29. Inouye M, Abraham G, Nelson CP, et al. Genomic Risk Prediction of Coronary Artery Disease in 480,000 Adults: Implications for Primary Prevention. J Am Coll Cardiol 2018;72:1883–93.

30. Oliynyk RT. Age-related late-onset disease heritability patterns and implications for genome-wide association studies. PeerJ 2019;7:e7168.

