## Supplementary material for "Polygenic and clinical risk scores and their impact on age at onset of cardiometabolic diseases and common cancers"

### FinnGen

#### Steering Committee

| | |
|---|---|
| Aarno Palotie | University of Helsinki / FIMM |
| Mark Daly | University of Helsinki / FIMM |
| **Pharma** | |
| Howard Jacob | Abbvie |
| Athena Matakidou | Astra Zeneca |
| Heiko Runz | Biogen |
| Sally John | Biogen |
| Robert Plenge | Celgene |
| Julie Hunkapiller | Genentech |
| Meg Ehm | GSK |
| Dawn Waterworth | GSK |
| Caroline Fox | Merck |
| Anders Malarstig | Pfizer |
| Kathy Klinger | Sanofi |
| Kathy Call | Sanofi |
| **UH & Biobanks** | |
| Tomi Mäkelä | University of Helsinki / FIMM |
| Jaakko Kaprio | University of Helsinki / FIMM |
| Petri Virolainen | Auria BB / Univ. of Turku /VSSHP |
| Kari Pulkki | Auria BB / Univ. of Turku /VSSHP |
| Terhi Kilpi | THL Biobank (BB) / THL |
| Markus Perola | THL Biobank (BB) / THL |
| Jukka Partanen | Finnish Red Cross Blood Service/FHRB |
| Anne Pitkäranta | HUS/Univ Hosp Districts |
| Riitta Kaarteenaho | Borealis BB/Univ. of Oulu/PPSHP |
| Seppo Vainio | Borealis BB/Univ. of Oulu/PPSHP |
| Kimmo Savinainen | Tampere BB/Univ Tampere/PSHP |
| Veli-Matti Kosma | Eastern Finland BB/UEF/PSSHP |
| Urho Kujala | Central Finland BB /University of Jyväskylä |
| **Other Experts/ Non-Voting Members** | |
| Outi Tuovila | Business Finland |
| Minna Hendolin | Business Finland |
| Raimo Pakkanen | Business Finland |

#### Scientific Committee

| | |
|---|---|
| **Pharma** | |
| Jeff Waring | Abbvie |
| Bridget Riley-Gillis | AbbVie |
| Athena Matakidou | Astra Zeneca |
| Heiko Runz | Biogen |
| Jimmy Liu | Biogen |
| Shameek Biswas | Celgene |
| Julie Hunkapiller | Genentech |
| Dawn Waterworth | GSK |
| Meg Ehm | GSK |
| Josh Hoffman | GSK |
| Dorothee Diogo | Merck |
| Caroline Fox | Merck |
| Anders Malarstig | Pfizer |
| Catherine Marshall | Pfizer |
| Xinli Hu | Pfizer |
| Kathy Call | Sanofi |
| Kathy Klinger | Sanofi |
| **UH & Biobanks** | |
| Samuli Ripatti | University of Helsinki / FIMM |
| Johanna Schleutker | Auria BB / Univ. of Turku /VSSHP |
| Markus Perola | THL Biobank (BB) / THL |
| Tiina Wahlfors | Finnish Red Cross Blood Service/FHRB |
| Olli Carpen | HUS/Univ Hosp Districts |
| Johanna Myllyharju | Borealis BB/Univ. of Oulu/PPSHP |
| Johannes Kettunen | Borealis BB/Univ. of Oulu/PPSHP |
| Reijo Laaksonen | Tampere BB/Univ Tampere/PSHP |
| Arto Mannermaa | Eastern Finland BB/UEF/PSSHP |
| Juha Paloneva | Central Finland BB /University of Jyväskylä/KSSHP |
| Urho Kujala | Central Finland BB /University of Jyväskylä |
| **Other Experts/ Non-Voting Members** | |
| Outi Tuovila | Business Finland |
| Minna Hendolin | Business Finland |
| Raimo Pakkanen | Business Finland |

### Clinical Groups

**Neurology Group**

| | |
|---|---|
| Hilkka Soininen LEAD | Kuopio |
| Valtteri Julkunen | Kuopio |
| Anne Remes | Oulu |
| Reetta Kälviäinen | Kuopio |
| Mikko Hiltunen | Kuopio |
| Jukka Peltola | Tampere |
| Pentti Tienari | Helsinki |
| Juha Rinne | Turku |
| Adam Ziemann | AbbVie |
| Jeffrey Waring | AbbVie |
| Sahar Esmaeeli | AbbVie |
| Nizar Smaoui | AbbVie |
| Anne Lehtonen | AbbVie |
| Susan Eaton | Biogen |
| Heiko Runz | Biogen |
| Sanni Lahdenperä | Biogen |
| Janet van Adelsberg | Celgene |
| Shameek Biswas | Celgene |
| John Michon | Genentech |
| Geoff Kerchner | Genentech |
| Julie Hunkapiller | Genentech |
| Natalie Bowers | Genentech |
| Edmond Teng | Genentech |
| John Eicher | Merck |
| Vinay Mehta | Merck |
| Padhraig Gormley | Merck |
| Kari Linden | Pfizer |
| Christopher Whelan | Pfizer |
| Fanli Xu | GSK |
| David Pulford | GSK |

**Gastroenterology Group**

| | |
|---|---|
| Martti Färkkilä LEAD | Helsinki |
| Sampsa Pikkarainen | HUS |
| Airi Jussila | Tampere |
| Timo Blomster | Oulu |
| Mikko Kiviniemi | Kuopio |
| Markku Voutilainen | Turku |
| Bob Georgantas | AbbVie |
| Graham Heap | AbbVie |
| Jeffrey Waring | AbbVie |
| Nizar Smaoui | AbbVie |
| Fedik Rahimov | AbbVie |
| Anne Lehtonen | AbbVie |
| Keith Usiskin | Celgene |
| Tim Lu | Genentech |
| Natalie Bowers | Genentech |
| Danny Oh | Genentech |
| John Michon | Genentech |
| Vinay Mehta | Merck |
| Dermot Reilly | Merck |
| Kirsi Kalpala | Pfizer |
| Melissa Miller | Pfizer |
| Xinli Hu | Pfizer |
| Linda McCarthy | GSK |

**Rheumatology Group**

| | |
|---|---|
| Kari Eklund LEAD | Helsinki |
| Antti Palomäki | Turku |
| Pia Isomaki | Tampere |
| Laura Pirilä | Turku |
| Oili Kaipiainen-Seppänen | Kuopio |
| Tuulikki Sokka-Isler | KSSHP |
| Markku Kauppi | Päijät-Häme Central Hospital/University of Tampere |
| Johanna Huhtakangas | Oulu |
| Eleanore Wigmore | AstraZeneca |
| Bob Georgantas | AbbVie |
| Jeffrey Waring | AbbVie |
| Fedik Rahimov | AbbVie |
| Apinya Lertratanakul | AbbVie |
| Nizar Smaoui | AbbVie |
| Anne Lehtonen | AbbVie |
| Marla Hochfeld | Celgene |
| Natalie Bowers | Genentech |
| John Michon | Genentech |
| Dorothee Diogo | Merck |
| Vinay Mehta | Merck |
| Kirsi Kalpala | Pfizer |
| Nan Bing | Pfizer |
| Xinli Hu | Pfizer |
| Jorge Esparza Gordillo | GSK |
| Nina Mars | University of Helsinki / FIMM |

**Pulmonology Group**

| | |
|---|---|
| Tarja Laitine LEAD | Tampere |
| Margit Pelkonen | Kuopio |
| Paula Kauppi | Helsinki |
| Hannu Kankaanranta | Tampere |
| Terttu Harju | Oulu |
| Nizar Smaoui | AbbVie |
| Eleanore Wigmore | AstraZeneca |
| Susan Eaton | Biogen |
| Steven Greenberg | Celgene |
| Hubert Chen | Genentech |
| Natalie Bowers | Genentech |
| John Michon | Genentech |
| Vinay Mehta | Merck |
| Jo Betts | GSK |
| Soumitra Ghosh | GSK |

**Cardiometabolic Diseases Group**

| | |
|---|---|
| Veikko Salomaa LEAD | THL |
| Teemu Niiranen | THL |
| Markus Juonala | Turku |
| Kaj Metsärinne | Turku |
| Mika Kähönen | Tampere |
| Juhani Junttila | Oulu |
| Markku Laakso | Kuopio |

| | |
|---|---|
| Jussi Pihlajamäki | Kuopio |
| Juha Sinisalo | Helsinki |
| Marja-Riitta Taskinen | Helsinki |
| Tiinamaija Tuomi | Helsinki |
| Jari Laukkanen | Keski-Suomen Keskussairaala/ University of Jyväskylä |
| Ben Challis | AstraZeneca |
| Keith Usiskin | Celgene |
| Andrew Peterson | Genentech |
| Julie Hunkapiller | Genentech |
| Natalie Bowers | Genentech |
| John Michon | Genentech |
| Dorothee Diogo | Merck |
| Dermot Reilly | Merck |
| Audrey Chu | Merck |
| Vinay Mehta | Merck |
| Jaakko Parkkinen | Pfizer |
| Melissa Miller | Pfizer |
| Anthony Muslin | Sanofi |
| Dawn Waterworth | GSK |

**Oncology Group**

| | |
|---|---|
| Heikki Joensuu LEAD | Helsinki |
| Tuomo Meretoja | Helsinki |
| Olli Carpen | Helsinki |
| Lauri Aaltonen | Helsinki |
| Annika Auranen | Tampere |
| Peeter Karihtala | Oulu |
| Saila Kauppila | Oulu |
| Päivi Auvinen | Kuopio |
| Klaus Elenius | Turku |
| Relja Popovic | AbbVie |
| Jeffrey Waring | AbbVie |
| Bridget Riley-Gillis | AbbVie |
| Anne Lehtonen | AbbVie |
| Athena Matakidou | AstraZeneca |
| Jennifer Schutzman | Genentech |
| Julie Hunkapiller | Genentech |
| Natalie Bowers | Genentech |
| John Michon | Genentech |
| Vinay Mehta | Merck |
| Andrey Loboda | Merck |
| Aparna Chhibber | Merck |
| Heli Lehtonen | Pfizer |
| Stefan McDonough | Pfizer |
| Marika Crohns | Sanofi |
| Diptee Kulkarni | GSK |

**Opthalmology Group**

| | |
|---|---|
| Kai Kaarniranta LEAD | Kuopio |
| Joni Turunen | HUS/ Secretary |
| Terhi Ollila | HUS |
| Sanna Seitsonen | HUS |
| Hannu Uusitalo | Tampere |
| Vesa Aaltonen | Turku |
| Hannele Uusitalo-Järvinen | PSHP |
| Marja Luodonpää | Oulu |
| Nina Hautala | Oulu |
| Heiko Runz | Biogen |
| Stephanie Loomis | Biogen |
| Erich Strauss | Genentech |
| Natalie Bowers | Genentech |
| Hao Chen | Genentech |
| John Michon | Genentech |
| Anna Podgornaia | Merck |
| Vinay Mehta | Merck |
| Dorothee Diogo | Merck |
| Joshua Hoffman | GSK |

**Dermatology Group**

| | |
|---|---|
| Kaisa Tasanen LEAD | Oulu |
| Laura Huilaja | Oulu |
| Katariina Hannula-Jouppi | HUS |
| Teea Salmi | Tampere |
| Sirkku Peltonen | Turku |
| Leena Koulu | Turku |
| Ilkka Harvima | Kuopio |
| Kirsi Kalpala | Pfizer |
| Ying Wu | Pfizer |
| David Choy | Genentech |
| John Michon | Genentech |
| Nizar Smaoui | AbbVie |
| Fedik Rahimov | AbbVie |
| Anne Lehtonen | AbbVie |
| Dawn Waterworth | GSK |

### FinnGen Teams

**Administration Team**

| | |
|---|---|
| Anu Jalanko | University of Helsinki / FIMM |
| Risto Kajanne | University of Helsinki / FIMM |
| Ulrike Lyhs | University of Helsinki / FIMM |

**Communication**

| | |
|---|---|
| Mari Kaunisto | University of Helsinki / FIMM |

**Analysis Team**

| | |
|---|---|
| Justin Wade Davis | Abbvie |
| Bridget Riley-Gillis | Abbvie |
| Danjuma Quarless | Abbvie |
| Slavé Petrovski | Astra Zeneca |
| Jimmy Liu | Biogen |
| Stephanie Loomis | Biogen |
| Paola Bronson | Biogen |
| Robert Yang | Celgene |
| Joseph Maranville | Celgene |
| Shameek Biswas | Celgene |

| | |
|---|---|
| Diana Chang | Genentech |
| Julie Hunkapiller | Genentech |
| Tushar Bhangale | Genentech |
| Natalie Bowers | Genentech |
| Dorothee Diogo | Merck |
| Emily Holzinger | Merck |
| Padhraig Gormley | Merck |
| Xulong Wang | Merck |
| Xing Chen | Pfizer |
| Åsa Hedman | Pfizer |
| Joshua Hoffman | GSK |
| Clarence Wang | Sanofi |
| Ethan Xu | Sanofi |
| Franck Auge | Sanofi |
| Clement Chatelain | Sanofi |
| Mitja Kurki | University of Helsinki / FIMM/ Broad Institute |
| Samuli Ripatti | University of Helsinki / FIMM |
| Mark Daly | University of Helsinki / FIMM |
| Juha Karjalainen | University of Helsinki / FIMM/ Broad Institute |
| Aki Havulinna | University of Helsinki / FIMM |
| Anu Jalanko | University of Helsinki / FIMM |
| Kimmo Palin | University of Helsinki |
| Priit Palta | University of Helsinki / FIMM |
| Pietro della Briotta Parolo | University of Helsinki / FIMM |
| Wei Zhou | Broad Institute |
| Susanna Lemmelä | University of Helsinki / FIMM |
| Manuel Rivas | University of Stanford |
| Jarmo Harju | University of Helsinki / FIMM |
| Aarno Palotie | University of Helsinki / FIMM |
| Arto Lehisto | University of Helsinki / FIMM |
| Andrea Ganna | University of Helsinki / FIMM |
| Vincent Llorens | University of Helsinki / FIMM |
| Antti Karlsson | Auria BB / Univ. of Turku /VSSHP |
| Kati Kristiansson | THL BB / THL |
| Mikko Arvas | Finnish Red Cross Blood Service BB /FHRB |
| Kati Hyvärinen | Finnish Red Cross Blood Service BB /FHRB |
| Jarmo Ritari | Finnish Red Cross Blood Service BB /FHRB |
| Tiina Wahlfors | Finnish Red Cross Blood Service BB /FHRB |
| Miika Koskinen | Helsinki BB/HUS/Univ Hosp Districts |
| Olli Carpen | Helsinki BB/HUS/Univ Hosp Districts |
| Johannes Kettunen | Borealis BB/Univ. of Oulu/PPSHP |
| Katri Pylkäs | Borealis BB/Univ. of Oulu/PPSHP |
| Marita Kalaoja | Borealis BB/Univ. of Oulu/PPSHP |
| Minna Karjalainen | Borealis BB/Univ. of Oulu/PPSHP |
| Tuomo Mantere | Borealis BB/Univ. of Oulu/PPSHP |
| Eeva Kangasniemi | Tampere BB/Univ Tampere/PSHP |
| Sami Heikkinen | Eastern Finland BB/UEFPSSHP |
| Arto Mannermaa | Eastern Finland BB/UEF/PSSHP |
| Eija Laakkonen | Central Finland BB /University of Jyväskylä/KSSHP |
| Juha Kononen | Central Finland BB /University of Jyväskylä/KSSHP |

**Sample Collection Coordination**

| | |
|---|---|
| Anu Loukola | Helsinki BB/HUS/Univ Hosp Districts |

**Sample Logistics**

| | |
|---|---|
| Päivi Laiho | THL BB / THL |
| Tuuli Sistonen | THL BB / THL |
| Essi Kaiharju | THL BB / THL |
| Markku Laukkanen | THL BB / THL |
| Elina Järvensivu | THL BB / THL |
| Sini Lähteenmäki | THL BB / THL |
| Lotta Männikkö | THL BB / THL |
| Regis Wong | THL BB / THL |

**Registry Data Operations**

| | |
|---|---|
| Kati Kristiansson | THL BB / THL |
| Hannele Mattsson | THL BB / THL |
| Susanna Lemmelä | University of Helsinki / FIMM |
| Tero Hiekkalinna | THL BB / THL |
| Manuel González Jiménez | THL BB / THL |

**Genotyping**

| | |
|---|---|
| Kati Donner | University of Helsinki / FIMM |

**Sequencing Informatics**

| | |
|---|---|
| Priit Palta | University of Helsinki / FIMM |

| | |
|---|---|
| Kalle Pärn | University of Helsinki / FIMM |
| Javier Nunez-Fontarnau | University of Helsinki / FIMM |

**Data Management and IT Infrastructure**

| | |
|---|---|
| Jarmo Harju | University of Helsinki / FIMM |
| Elina Kilpeläinen | University of Helsinki / FIMM |
| Timo P. Sipilä | University of Helsinki / FIMM |
| Georg Brein | University of Helsinki / FIMM |
| Alexander Dada | University of Helsinki / FIMM |
| Ghazal Awaisa | University of Helsinki / FIMM |
| Anastasia Shcherban | University of Helsinki / FIMM |
| Tuomas Sipilä | University of Helsinki / FIMM |

**Clinical Endpoint Development**

| | |
|---|---|
| Hannele Laivuori | University of Helsinki / FIMM |
| Aki Havulinna | University of Helsinki / FIMM |
| Susanna Lemmelä | University of Helsinki / FIMM |
| Tuomo Kiiskinen | University of Helsinki / FIMM |

**Trajectory Team**

| | |
|---|---|
| Tarja Laitinen | Tampere University Hospital |
| Harri Siirtola | University of Tampere |
| Javier Gracia Tabuenca | University of Tampere |

**Biobank Directors**

| | |
|---|---|
| Lila Kallio | Auria Biobank |
| Sirpa Soini | THL Biobank |
| Jukka Partanen | Blood Service Biobank |
| Kimmo Pitkänen | Helsinki Biobank |
| Seppo Vainio | Northern Finland Biobank Borealis |
| Kimmo Savinainen | Tampere Biobank |
| Veli-Matti Kosma | Biobank of Eastern Finland |
| Teijo Kuopio | Central Finland Biobank |

### Genotyping and imputation in FinnGen

FinnGen samples were genotyped with Illumina and Affymetrix arrays (Illumina Inc., San Diego, and Thermo Fisher Scientific, Santa Clara, CA, USA) and put through the same rigorous QC steps as described above. Genotype imputation was carried out by using the population-specific SISu v3 imputation reference panel with Beagle 4.1 (version 08Jun17.d8b, https://faculty.washington.edu/browning/beagle/b4_1.html) as described in the following protocol: dx.doi.org/10.17504/protocols.io.nmndc5e. Post-imputation QC involved excluding variants with imputation INFO <0.7.

### Genotyping and imputation in FINRISK

26,404 FINRISK samples were genotyped using several arrays: the HumanCoreExome BeadChip, the Human610-Quad BeadChip, the Affymetrix6.0, and the Infinium HumanOmniExpress (Illumina Inc., San Diego and Affymetrix, Inc., Santa Clara, CA, USA). Genotype calls were generated together with other available data sets using zCall at the Institute for Molecular Medicine Finland (FIMM). After sample-wise quality control (exclude samples with ambiguous gender, missingness (>5%), excess heterozygosity (+-4SD), non-European ancestry) and variant-wise quality control (exclude SNPs with high missingness (>2%), low HWE P-value (<1e-6), minor allele count (MAC) <3 (in case Zcall'ed chip data) or MAC <10 (chip data called using Illumina GenCall) steps, the samples were pre-phased using Eagle2 (version 2.3). Genotype imputation was carried out by using a Finnish population-specific reference panel consisting of 2690 high-coverage WGS and 5092 WES samples with IMPUTE2 (version 2.3.2) that allows the usage of two panels at the same time (the 'merge_ref_panels' option). Post-imputation quality control involved excluding variants imputed with imputation INFO < 0.7. Chromosome X variants were also excluded from the downstream analyses. We excluded one individual of each sample-pair with kinship >0.125, and calculated principal components for the unrelated individuals. The 26,404 samples contained the 2012 FINRISK cohort; this study used only FINRISK cohorts from 1992, 1997, 2002, and 2007, comprising 21,813 unrelated individuals.

### Polygenic risk scores

In LDpred,[1] using a linkage disequilibrium (LD) reference panel matching the genome-wide association study (GWAS) discovery population is recommended. We applied the Finnish panel SISu v2, however, type 2 diabetes (T2D) PRSs calculated with i) SISuv2 and ii) 503 Europeans from the 1000 Genomes phase 3[2] as the LD reference showed high correlation (>0.9). Due to the high LD in the isolated Finnish population, we selected an LD-radius approximately twice the radius recommended, which is M/3,000, where M is the total number of single nucleotide polymorphisms used in the analysis. Variants with minor allele frequency less than 1% are excluded by the software.

We calculated the polygenic risk scores by summing the dosage of each risk allele carried by an individual (ranging from 0 to 2 for each variant, dosage used for incorporating imputation uncertainty), weighting each variant by its natural logarithm of the relative risk extracted from the genome-wide association study. For each individual *i*, this results in a single value on a continuous scale:

$$\text{PRS}_i = \sum_{j=1}^{M} \hat{\beta}_j \times dosage_{ij}$$

where $\hat{\beta}_j$ is the weight for variant *j* obtained from GWAS summary statistics. Within the whole FINRISK dataset, the PRS the highest C-index was chosen for the subsequent analyses (Supplementary Table 4). Adjusted survival curves were plotted with the R package *survminer*, using the calculation parameter "conditional", which after rebalancing averages for the polygenic risk score categories. The restricted mean survival times[3] (RMST; age 85 as the upper limit) were estimated by fitting flexible parametric survival models, which generated similar effect sizes as the Cox proportional hazards models.

### Clinical risk calculators

In the FINRISK analyses, the association between PRS and disease was tested for incident cases only. The number of prevalent cases excluded was 954 for coronary heart disease, 671 for type 2 diabetes (T2D), 351 for atrial fibrillation (AF), 164 for breast cancer, and 59 for prostate cancer. For clinical risk factor variables missingness was at most 1.8%; individuals with missing data were removed from each respective clinical risk assessment. For calculating the 10-year risk for hard atherosclerotic cardiovascular disease (ASCVD) according to the Pooled Cohort Equations by ACC/AHA (2013), were excluded 23 individuals with missing data in any of the risk factors.

Clinical high risk for T2D was defined by the criteria for testing diabetes or prediabetes in asymptomatic adults, based on American Diabetes Association's current clinical practice recommendations.[4] These criteria consist of a combination of BMI ≥25kg/m$^2$ and one or more additional risk factors, of which the following were applicable in our study: first-degree relative with diabetes, history of cardiovascular disease, hypertension (≥140/90 mmHg), HDL <0.90 mmol/l, triglyceride level >2.82 mmol/l, and severe obesity (BMI ≥35kg/m$^2$). Individuals who with impaired fasting glucose (≥5.6 mmol/L) were also defined as having high risk. In assessing this risk, history of cardiovascular disease was defined as physician-diagnosed coronary heart disease or stroke (see Supplementary Table 2 for definition of coronary heart disease; stroke was any of I61, I63, I64 except I63.6 (International Classification of Diseases, 10$^{th}$ revision; ICD-10) or 431, 4330A, 4331A, 4339A, 4340A, 4341A, 4349A, 436 (ICD-9) as the underlying or direct cause of death, or as the main or side diagnosis at hospital discharge. 82 individuals with missing data on BMI were excluded from these analyses involving clinical risk assessment of T2D.

When taking all calculator components from the original study, the original CHARGE-AF score showed poor calibration with a mean 5-year risk 0.02% in individuals aged ≥45. To improve calibration, we obtained the mean component from FINRISK individuals aged 45 to 74, which resulted in a 5-year mean risk of 4.2% (standard deviation 4.3%). We did not revise the baseline hazard, as the original baseline hazard ≈0.972 was similar to ours (≈0.977). 85 individuals with missing data for the risk variables were excluded.

### Supplementary References

1. Vilhjalmsson BJ, Yang J, Finucane HK, et al. Modeling Linkage Disequilibrium Increases Accuracy of Polygenic Risk Scores. Am J Hum Genet 2015;97:576-92.
2. Genomes Project Consortium, Auton A, Brooks LD, et al. A global reference for human genetic variation. Nature 2015;526:68-74.
3. Royston P, Parmar MK. Restricted mean survival time: an alternative to the hazard ratio for the design and analysis of randomized trials with a time-to-event outcome. BMC Med Res Methodol 2013;13:152.
4. American Diabetes Association. 2. Classification and Diagnosis of Diabetes: Standards of Medical Care in Diabetes-2019. Diabetes Care 2019;42:S13-S28.
5. Zhou W, Nielsen JB, Fritsche LG, et al. Efficiently controlling for case-control imbalance and sample relatedness in large-scale genetic association studies. Nat Genet 2018;50:1335-41.

**Supplementary Figure 1.** Manhattan plot (A) and quantile-quantile plot (B) for the genome-wide association study for PheCode 411 (Ischemic heart disease) by Zhou et al.[5] These summary statistics were used for constructing the polygenic risk score for coronary heart disease.

**A**

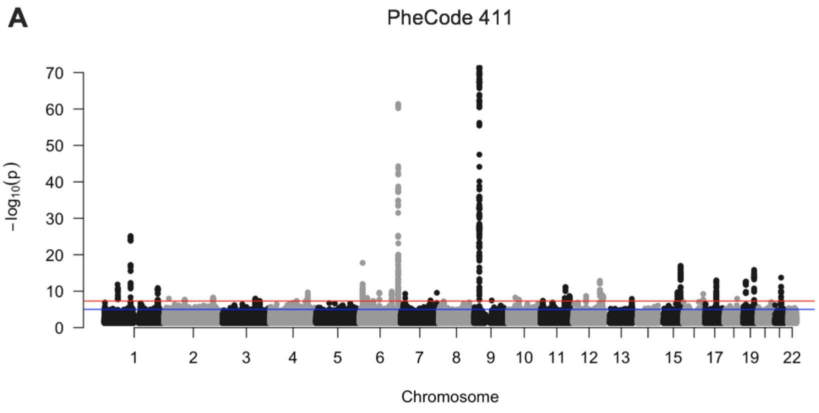

**B**

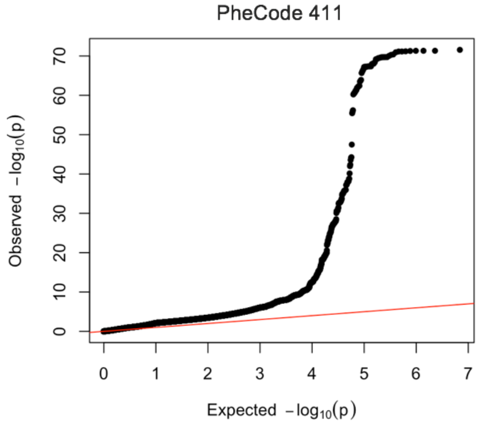

**Supplementary Figure 2.** With any breast cancer as the outcome in FinnGen, adjusted survival curves for estrogen receptor-specific polygenic risk scores (PRS). The PRS for any breast cancer showed high correlation with the estrogen receptor-positive PRS (r = 0.93) and moderate correlation with estrogen receptor-negative PRS (0.54).

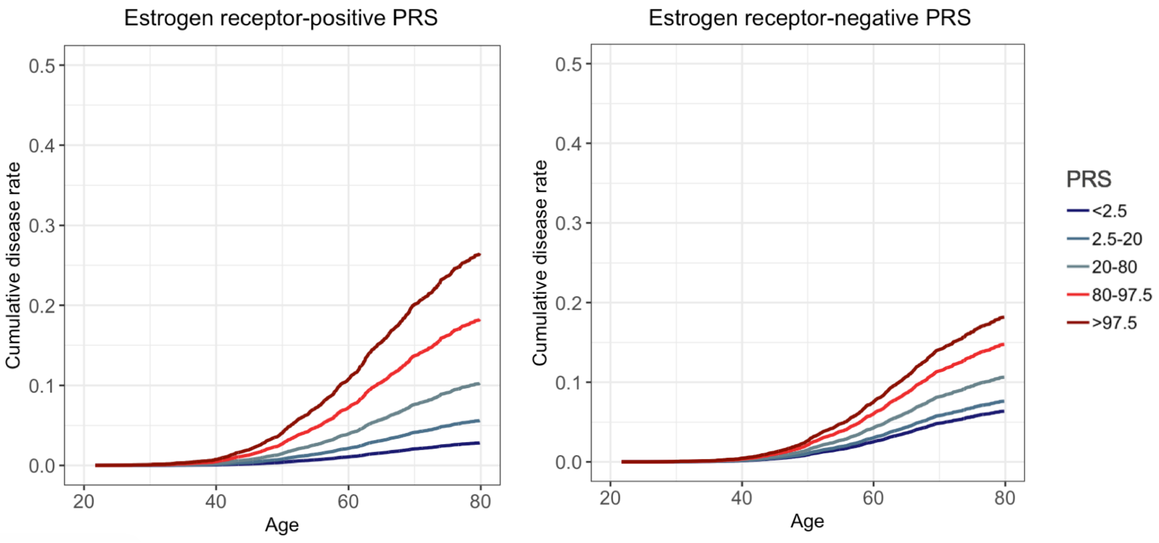

**Supplementary Figure 3.** Difference in age at disease onset by sex, across polygenic risk score categories (FinnGen).

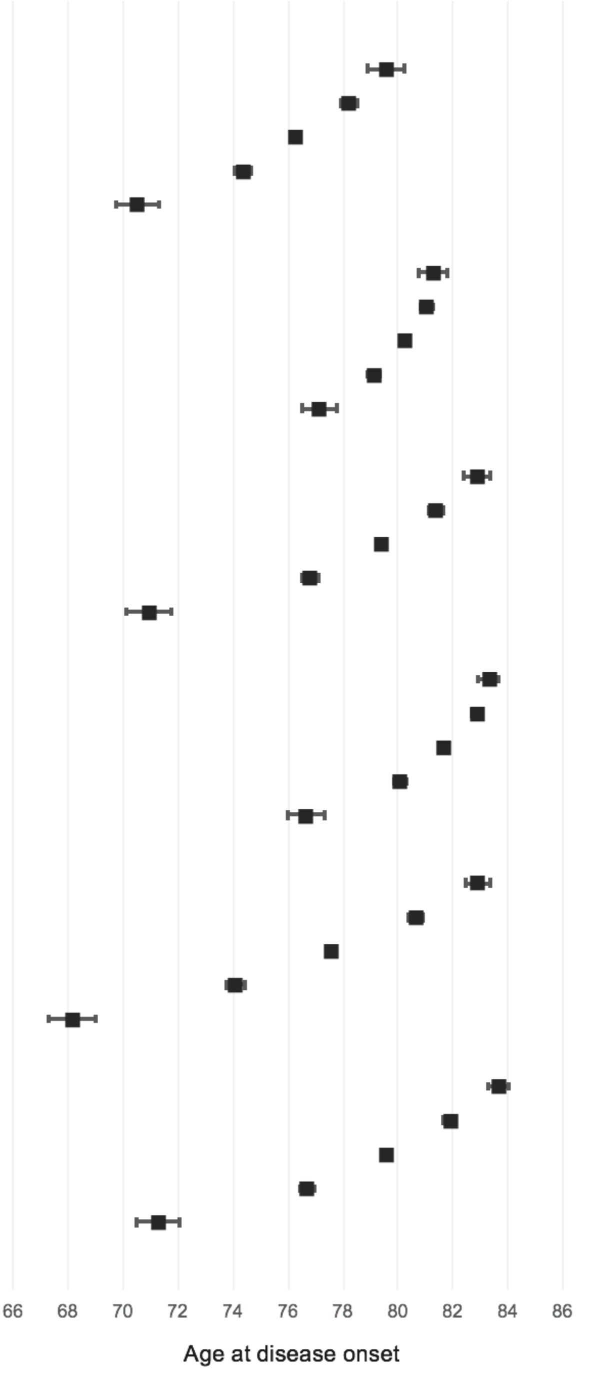

| | Mean difference in years (95% CI) | p |
|---|---|---|
| **CHD, men** | | |
| <2.5 | 3.30 (2.66, 3.94) | 6.27x10$^{-17}$ |
| 2.5-20 | 1.97 (1.71, 2.23) | 1.11x10$^{-43}$ |
| 20-80 | 0 | - |
| 80-97.5 | -1.89 (-2.15, -1.63) | 1.34x10$^{-50}$ |
| >97.5 | -5.73 (-6.48, -4.99) | 2.66x10$^{-66}$ |
| **CHD, women** | | |
| <2.5 | 1.04 (0.51, 1.58) | 6.04x10$^{-4}$ |
| 2.5-20 | 0.82 (0.62, 1.02) | 4.35x10$^{-14}$ |
| 20-80 | 0 | - |
| 80-97.5 | -1.10 (-1.30, -0.89) | 2.07x10$^{-29}$ |
| >97.5 | -3.08 (-3.70, -2.46) | 1.43x10$^{-31}$ |
| **AF, men** | | |
| <2.5 | 3.48 (3.00, 3.96) | 9.03x10$^{-20}$ |
| 2.5-20 | 2.01 (1.79, 2.24) | 1.46x10$^{-49}$ |
| 20-80 | 0 | - |
| 80-97.5 | -2.56 (-2.82, -2.31) | <1.00x10$^{-100}$ |
| >97.5 | -8.44 (-9.21, -7.66) | <1.00x10$^{-100}$ |
| **AF, women** | | |
| <2.5 | 1.65 (1.26, 2.03) | 1.28x10$^{-9}$ |
| 2.5-20 | 1.21 (1.05, 1.38) | 1.72x10$^{-33}$ |
| 20-80 | 0 | - |
| 80-97.5 | -1.59 (-1.79, -1.39) | 2.33x10$^{-72}$ |
| >97.5 | -5.02 (-5.67, -4.37) | 7.99x10$^{-91}$ |
| **T2D, men** | | |
| <2.5 | 5.33 (4.89, 5.77) | 2.35x10$^{-38}$ |
| 2.5-20 | 3.08 (2.83, 3.32) | 2.05x10$^{-92}$ |
| 20-80 | 0 | - |
| 80-97.5 | -3.49 (-3.78, -3.20) | <1.00x10$^{-100}$ |
| >97.5 | -9.42 (-10.25, -8.59) | <1.00x10$^{-100}$ |
| **T2D, women** | | |
| <2.5 | 4.08 (3.71, 4.44) | 3.82x10$^{-28}$ |
| 2.5-20 | 2.30 (2.09, 2.51) | 2.06x10$^{-69}$ |
| 20-80 | 0 | - |
| 80-97.5 | -2.88 (-3.13, -2.63) | <1.00x10$^{-100}$ |
| >97.5 | -8.29 (-9.07, -7.51) | <1.00x10$^{-100}$ |

CHD = coronary heart disease, AF = atrial fibrillation or flutter, T2D = type 2 diabetes. The estimands for age at onset are restricted mean survival times (RMST).

**Supplementary Figure 4.** Correlation between polygenic and clinical risk in FINRISK, using respective incident disease cases and controls.

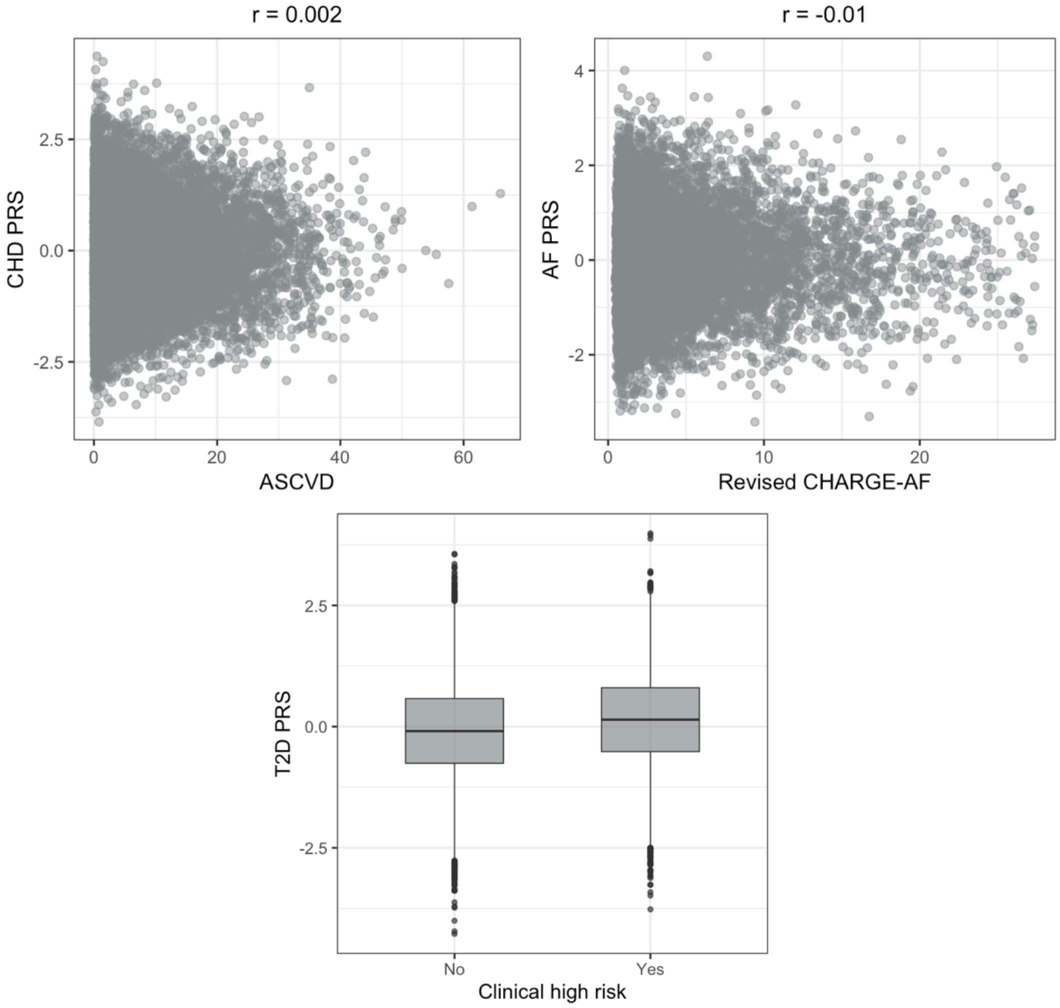

CHD = coronary heart disease, PRS = polygenic risk score, AF = atrial fibrillation or flutter, T2D = type 2 diabetes, BMI = body mass index, WHR = waist-hip ratio. ASCVD = the 10-year risk calculator for hard atherosclerotic cardiovascular disease (ASCVD), according to the Pooled Cohort Equations by ACC/AHA (2013). Revised CHARGE-AF = revised version of the CHARGE-AF, a risk calculator for 5-year risk of AF. Clinical high risk for T2D defined as BMI ≥25kg/m2 and one of more additional clinical risk factors.[4]

**Supplementary Figure 5.** The area under the receiver operating characteristic curves (AUC) for high PRS, high clinical risk, or their combination.

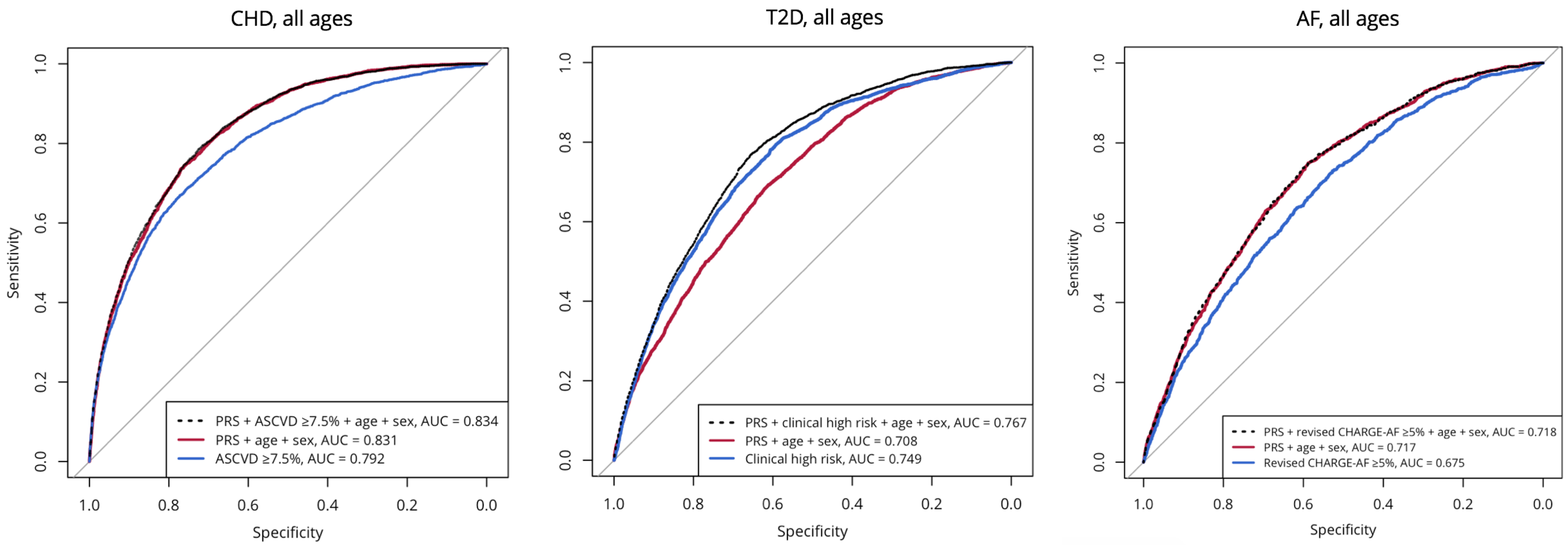

CHD = coronary heart disease (N = 20,165), T2D = type 2 diabetes, AF = atrial fibrillation or flutter (N = 10,666). High PRS = above the 90th percentile. ASCVD = the 10-year risk calculator for hard atherosclerotic cardiovascular disease (ASCVD), according to the Pooled Cohort Equations by ACC/AHA (2013). Clinical high risk for T2D defined as BMI ≥25kg/m$^2$ and one of more additional clinical risk factors.[16] Revised CHARGE-AF = revised version of the CHARGE-AF, a risk calculator for the 5-year risk of AF.

**Supplementary Figure 6.** Adjusted survival curves in FINRISK, showing cumulative risk of incident disease in by polygenic risk score (PRS) categories.

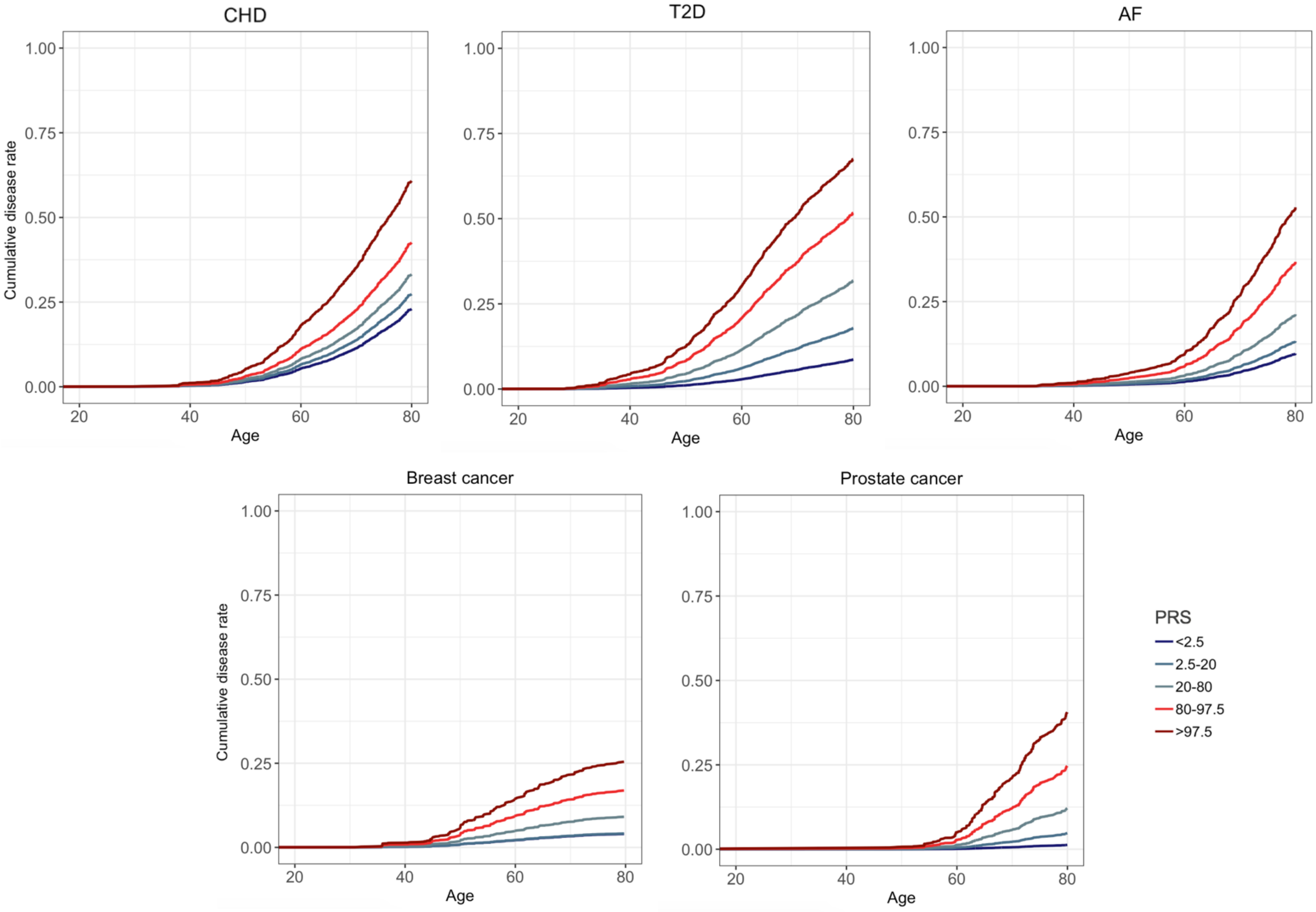

The FINRISK cohorts (n = 21,813) comprised of 2,197 incident cases of CHD, 1,431 cases of AF, 2,516 cases of T2D, 404 cases of breast cancer, and 444 cases of prostate cancer.

**Table S1.** The prospective epidemiological and disease-based cohorts, and hospital biobank samples in FinnGen Data Freeze 2.

| **Cohort** | **N** |
|---|---|
| Auria biobank* | 9,967 |
| Blood Service biobank | 1,3222 |
| Borealis biobank* | 1,368 |
| Botnia Family | 1,216 |
| Botnia New | 6 |
| Botnia PPP | 4,856 |
| Botnia Sib-Helsinki | 431 |
| Corogene | 4,495 |
| Eastern Finland biobank* | 1,965 |
| FinHealth 2017 | 5,783 |
| FINRISK 1992-2012 | 29,550 |
| GeneRISK | 6,960 |
| Health 2000 | 6,602 |
| Health 2011 | 711 |
| Helsinki biobank* | 21,014 |
| Kuusamo 2011 | 145 |
| Migraine | 7,732 |
| SUPER | 4,402 |
| Tampere biobank* | 1,973 |
| THL Diabetes | 6,983 |
| Twins | 5,919 |
| **Sum** | **135,300** |

*Hospital-based biobanks

**Table S2.** Disease endpoint definitions.

| | Additional definitions | Only main diagnosis accepted | ICD-10 | ICD-9 | ICD-8 | ICD-10 exclu-sions | Cause of death ICD-10 | Cause of death ICD-9 | Cause of death ICD-8 | Cause of death ICD-10 exclu-sions | Cause of death ICD-9 exclu-sions | Topo-graphical codes* |
|---|---|---|---|---|---|---|---|---|---|---|---|---|
| **Coronary heart disease** | Myocardial infarction\|Myocardial infarction, strict\|Complications following myocardial infarction\|Prior myocardial infactrion\|Angina pectoris\|Other coronary atherosclerosis\|Coronary artery bypass graft**\|Coronary angioplasty** | | | | | | | | | | | |
| **Major coronary heart disease event** | Myocardial infarction\|Coronary artery bypass graft\|Coronary angioplasty | Yes | I20.0 \| I21 \| I22 | 410 \| 411.0 | 410 \| 411.0 | | I2[1-5] \| I46 \| R96 \| R98 | 41[0-4] \| 798 | 41[0-4] \| 798 | | 798.0A | |
| **Myocardial infarction, strict** | | Yes | I21 \| I22 | 410 | 410 | | I21 \| I22 | 410 | 410 | | | |
| **Myocardial infarction** | | | I21 \| I22 | 410 | 410 | | I21 \| I22 | 410 | 410 | | | |
| **Complications following myocardial infarction** | | | I23 | - | - | | I23 | - | - | | | |
| **Old myocardial infarction** | | | I25.2 | 412 | 412 | | I25.3 | 412 | 412 | | | |
| **Angina pectoris** | | | I20 | 413 \| 411[0-1] | 413 | | I20 | 413 \| 411[0-1] | 413 | | | |
| **Other coronary atheroclerosis** | | | I25 \| I24 \| Z95.1 \| T82.2 | 414 \| 996.0A | 414 | I25.3 | I25 \| I24 \| Z95.1 \| T82.2 | 414 \| 996.0A | 414 | I25.3 | | |

| | | | | | | | | | |
|---|---|---|---|---|---|---|---|---|---|
| **Atrial fibrillation and flutter** | Eligibility for special reimbursement for apixaban, dabigatran, edoxaban, rivaroxaban or dronedarone for ICD-10 I48 | I48 | 427.3 | 427.92 | | I48 | 427.3 | 427.92 | |
| **Malignant neoplasm of breast / breast cancer** | | | | | | | | | C50 |
| **Malignant neoplasm of prostate / prostate cancer** | | | | | | | | | C61 |
| **Type 2 diabetes*** ** | Any type 2 diabetes diagnosis defined below \| Medication purchases for ATC A10B, Blood glucose lowering drugs, excluding insulins. | | | | E10[0-9] | | | | |
| Type 2 diabetes with coma | | E11.0 | 250.2A | - | | E11.0 | 250.2A | | |
| Type 2 diabetes with ketoacidosis | | E11.1 | 250.1A | - | | E11.1 | 250.1A | | |
| Type 2 diabetes with renal complications | | E11.2 | 250.3A | - | | E11.2 | 250.3A | | |
| Type 2 diabetes with ophthalmic complications | | E11.3 | 250.4A | - | | E11.3 | 250.4A | | |
| Type 2 diabetes with neurological complications | | E11.4 | 250.5A | - | | E11.4 | 250.5A | | |
| Type 2 diabetes with peripheral circulatory complications | | E11.5 | 250.6A | - | | E11.5 | 250.6A | | |
| Type 2 diabetes with other specified/ multiple/unspecifie d complications | Eligibility for medication reimbursement with ICD-10 E11 | E11[6-8] | 250.7A \| 250.8A | - | | E11[6-8] | 250.7A \| 250.8A | | |
| Type 2 diabetes without complications | | E11.9 | 250.0A | - | | E11.9 | 250.0A | | |

* The International Classification of Diseases for Oncology, Third Edition (ICD-O-3). **Procedure code identified at hospital discharge or from the nationwide register of invasive cardiac procedures. *** In FinnGen analyses, individuals with type 1 diabetes were excluded from cases (ICD-10 E10[0-9], ICD-9 250[0-8]B as a hospital discharge diagnosis or cause of death, or E10 for medication reimbursement)

**Table S3.** Genome-wide association studies used for constructing the polygenic risk scores and the number of variants in the final scores.

| | GWAS summary statistics source | Article link | Data download link | Most recent access to data download | SNPs in discovery GWAS | SNPs in PRS calculation | LD radius | Additional information |
|---|---|---|---|---|---|---|---|---|
| **Coronary heart disease** | UKBB SAIGE | https://www.nature.com/articles/s41588-018-0184-y | https://www.dropbox.com/sh/wuj4y8wsqjz78om/AAACfAJK54KtvnzSTAoaZTLma?dl=0 | Nov 2, 2018 | 28 345 446 | 6 412 950 | 4 000 | PheCode 411 Ischemic heart disease |
| **Type 2 diabetes** | Mahajan et al 2018 | https://www.nature.com/articles/s41588-018-0241-6 | http://www.diagram-consortium.org/downloads.html | Dec 21, 2018 | 23 465 133 | 6 437 380 | 4 000 | Not adjusted for BMI |
| **Atrial fibrillation and flutter** | Nielsen et al 2018 | https://www.nature.com/articles/s41588-018-0171-3 | http://csg.sph.umich.edu/willer/public/afib2018/ | Dec 21, 2018 | 34 740 187 | 6 171 733 | 4 000 | |
| **Breast cancer** | Michailidou et al 2017 | https://www.nature.com/articles/nature24284 | http://bcac.ccge.medschl.cam.ac.uk/bcacdata/oncoarray/gwas-icogs-and-oncoarray-summary-results/ | Dec 21, 2018 | 11 792 358 | 6 390 808 | 4 000 | |
| **Breast cancer, estrogen receptor-positive** | Michailidou et al 2017 | https://www.nature.com/articles/nature24284 | http://bcac.ccge.medschl.cam.ac.uk/bcacdata/oncoarray/gwas-icogs-and-oncoarray-summary-results/ | Dec 21, 2018 | 11 784 434 | 6 390 799 | 4 000 | |
| **Breast cancer, estrogen receptor-negative** | Michailidou et al 2017 | https://www.nature.com/articles/nature24284 | http://bcac.ccge.medschl.cam.ac.uk/bcacdata/oncoarray/gwas-icogs-and-oncoarray-summary-results/ | Dec 21, 2018 | 11 784 725 | 6 390 805 | 4 000 | |
| **Prostate cancer** | Schumacher et al 2018 | https://www.nature.com/articles/s41588-018-0142-8 | http://practical.icr.ac.uk/blog/?page_id=8088 | Dec 21, 2018 | 20 734 509 | 6 606 785 | 4 000 | |

**Table S4.** The LDpred algorithm uses a tuning parameter p for denoting the fraction of variants assumed to be causal for the disease. The PRS with the highest C-index (bolded) in FINRISK was chosen for the subsequent analyses.

| | Fraction of causal markers | C-index |
|---|---|---|
| **Coronary heart disease** | 0.0001* | 0.8163 |
| | 0.0003* | 0.8162 |
| | 0.001* | 0.8163 |
| | **0.003** | **0.8203** |
| | 0.01 | 0.8195 |
| | 0.03 | 0.8188 |
| | 0.1 | 0.8184 |
| | 0.3 | 0.8183 |
| | 1 | 0.8183 |
| | inf | 0.8183 |
| **Type 2 diabetes** | 0.0001* | 0.7022 |
| | 0.0003* | 0.7043 |
| | 0.001* | 0.7033 |
| | 0.003* | 0.7033 |
| | 0.01* | 0.7089 |
| | 0.03* | 0.7091 |
| | 0.1 | 0.7374 |
| | **0.3** | **0.7417** |
| | 1 | 0.7402 |
| | inf | 0.7398 |
| **Atrial fibrillation or flutter** | 0.0001* | 0.7912 |
| | 0.0003* | 0.7915 |
| | 0.001* | 0.7921 |
| | 0.003* | 0.7916 |
| | 0.01* | 0.7942 |
| | **0.03** | **0.8135** |
| | 0.1 | 0.8107 |
| | 0.3 | 0.8082 |
| | 1 | 0.8057 |
| | inf | 0.8055 |
| **Prostate cancer** | 0.0001* | 0.8076 |
| | 0.0003* | 0.8077 |
| | 0.001* | 0.8096 |
| | 0.003 | 0.8140 |
| | **0.01** | **0.8416** |
| | 0.03 | 0.8341 |
| | 0.1 | 0.8270 |
| | 0.3 | 0.8237 |
| | 1 | 0.8224 |
| | inf | 0.8223 |
| **Breast cancer** | 0.0001* | 0.6426 |
| | 0.0003* | 0.6403 |
| | 0.001* | 0.6454 |
| | 0.003* | 0.6490 |
| | 0.01* | 0.6422 |

| | | |
|---|---|---|
| | **0.03** | **0.7042** |
| | 0.1 | 0.6955 |
| | 0.3 | 0.6892 |
| | 1 | 0.6852 |
| | inf | 0.6853 |
| **Breast cancer, estrogen receptor-positive** | 0.0001* | 0.6404 |
| | 0.0003* | 0.6421 |
| | 0.001* | 0.6434 |
| | 0.003* | 0.6450 |
| | 0.01* | 0.6479 |
| | **0.03** | **0.6990** |
| | 0.1 | 0.6912 |
| | 0.3 | 0.6868 |
| | 1 | 0.6834 |
| | inf | 0.6833 |
| **Breast cancer, estrogen receptor-negative** | 0.0001* | 0.6403 |
| | 0.0003* | 0.6410 |
| | 0.001* | 0.6405 |
| | **0.003** | **0.6511** |
| | 0.01 | 0.6472 |
| | 0.03 | 0.6449 |
| | 0.1 | 0.6438 |
| | 0.3 | 0.6435 |
| | 1 | 0.6434 |
| | inf | 0.6432 |

*One or multiple chromosomes failed to converge.

**Table S5.** Baseline characteristics for FINRISK.

| | FINRISK 1992<br>N = 4,745 | FINRISK 1997<br>N = 6,733 | FINRISK 2002<br>N = 5,427 | FINRISK 2007<br>N = 4,908 |
|---|---|---|---|---|
| Follow-up in years, mean (SD) | 22.3 (4.3) | 17.5 (3.7) | 13.3 (2.0) | 8.7 (1.0) |
| Age, mean (SD) | 44.3 (11.4) | 48.2 (13.4) | 48.3 (13.1) | 51.1 (13.9) |
| Age ≤50, % | 65.9 | 54.9 | 52.4 | 45.2 |
| Women, % | 53.8 | 51.0 | 53.4 | 53.3 |
| Current smokers, % | 28.0 | 23.6 | 26.3 | 19.9 |
| TC, mean (SD) | 5.6 (1.1) | 5.5 (1.1) | 5.6 (1.1) | 5.3 (1.0) |
| LDL, mean (SD) | 3.5 (1.0) | 3.5 (0.9) | 3.4 (1.0) | 3.2 (0.9) |
| HDL, mean (SD) | 1.4 (0.3) | 1.4 (0.4) | 1.5 (0.4) | 1.4 (0.4) |
| TG, mean (SD) | 1.5 (1.1) | 1.5 (1.0) | 1.4 (1.0) | 1.4 (0.9) |
| SBP, mean (SD) | 135.3 (19.3) | 136.0 (19.9) | 135.2 (20.0) | 136.2 (20.3) |
| BMI, mean (SD) | 26.1(4.4) | 26.6 (4.5) | 26.9 (4.7) | 27.2 (4.9) |
| WHR, mean (SD) | 0.8 (0.1) | 0.9 (0.1) | 0.9 (0.1) | 0.9 (0.1) |
| Blood pressure-lowering treatment, % | 9.0 | 13.0 | 14.3 | 21.3 |
| Lipid-lowering treatment, % | 1.5 | 3.2 | 7.1 | 14.6 |
| Positive family history for any diabetes, % | N/A | 25.8 | 26.4 | 28.7 |
| Positive family history for early MI, % | 23.6 | 25.5 | 25.6 | 15.5 |
| Prevalent CHD, % | 3.3 | 5.3 | 5.0 | 6.4 |
| Prevalent MI, % | 0.8 | 1.3 | 1.0 | 1.7 |
| Prevalent AF, % | 0.9 | 1.8 | 1.7 | 2.9 |
| Prevalent T2D, % | 0.4 | 2.8 | 3.8 | 4.5 |
| Prevalent breast cancer in women, % | 1.0 | 1.2 | 1.3 | 2.3 |
| Prevalent prostate cancer in men, % | 0.0 | 0.6 | 0.5 | 1.2 |
| Incident CHD, % | 14.2 | 13.8 | 8.8 | 5.3 |
| Incident MI, % | 5.6 | 5.7 | 3.7 | 2.0 |
| Incident AF, % | 9.5 | 9.1 | 6.0 | 4.0 |
| Incident T2D, % | 15.7 | 13.2 | 10.1 | 7.2 |
| Incident breast cancer in women, % | 5.3 | 4.1 | 3.0 | 1.5 |
| Incident prostate cancer in men, % | 5.5 | 5.6 | 3.6 | 2.0 |

TC = total cholesterol, LDL = low-density lipoprotein (using the Friedewald equation), HDL = high-density lipoprotein, TG = triglycerides, SBP = systolic blood pressure, BMI = body mass index, WHR = waist-hip ratio, CHD = coronary heart disease, MI = myocardial infarction, T2D = type 2 diabetes, AF = atrial fibrillation or flutter. Units: lipid measurements mmol/l, SBP mmHg, BMI kg/m2.

**Table S6.** Impact of family history on polygenic risk score (PRS) effect size estimates (per standard deviation increment).

| | **HR (95% CI)** | **p** |
|---|---|---|
| CHD PRS | 1.27 (1.22-1.32) | $4.77 \times 10^{-28}$ |
| Family history of early MI | 1.49 (1.36-1.63) | $7.57 \times 10^{-18}$ |
| CHD PRS + family history of early MI | | |
| CHD PRS | 1.26 (1.21-1.31) | $5.24 \times 10^{-26}$ |
| Family history of early MI | 1.45 (1.32-1.59) | $1.21 \times 10^{-15}$ |
| T2D PRS | 1.57 (1.50-1.66) | $1.24 \times 10^{-68}$ |
| Family history of any diabetes | 1.62 (1.47-1.78) | $1.67 \times 10^{-22}$ |
| T2D PRS + family history of any diabetes | | |
| T2D PRS | 1.54 (1.46-1.62) | $2.24 \times 10^{-62}$ |
| Family history of early MI | 1.49 (1.35-1.64) | $8.27 \times 10^{-16}$ |

CHD = coronary heart disease, T2D = type 2 diabetes. T2D models adjusted for BMI.

**Table S7.** Hazard ratios (HR) and 95% confidence intervals (CI) per standard deviation increment in FINRISK.

| | **HR (95% CI)** | **p** |
|---|---|---|
| Coronary heart disease | 1.25 (1.18-1.32) | $1.74 \times 10^{-14}$ |
| Type 2 diabetes | 1.70 (1.63-1.78) | $<1.00 \times 10^{-100}$ |
| Atrial fibrillation or flutter | 1.62 (1.54-1.70) | $8.85 \times 10^{-78}$ |
| Breast cancer | 1.75 (1.59-1.92) | $2.61 \times 10^{-30}$ |
| Prostate cancer | 1.88 (1.71-2.06) | $4.74 \times 10^{-41}$ |

**Table S8.** Hazard ratios (HR) and 95% confidence intervals (CI) for polygenic risk score (PRS) bins in FINRISK.

| | HR (95% CI) | p | N cases / N contols |
|---|---|---|---|
| **CHD PRS** | | | |
| <2.5 | 0.65 (0.47-0.89) | 0.008 | 39 / 466 |
| 2.5-20 | 0.81 (0.72-0.92) | 0.001 | 313 / 3,220 |
| 20-80 | 1 (reference) | - | 1,274 / 10,838 |
| 80-97.5 | 1.35 (1.22-1.50) | $2.45 \times 10^{-8}$ | 471 / 3,062 |
| >97.5 | 2.42 (1.97-2.97) | $2.21 \times 10^{-17}$ | 100 / 405 |
| **T2D PRS** | | | |
| <2.5 | 0.23 (0.14-0.38) | $7.14 \times 10^{-9}$ | 16 / 513 |
| 2.5-20 | 0.50 (0.43-0.58) | $2.80 \times 10^{-21}$ | 224 / 3,476 |
| 20-80 | 1 (reference) | - | 1,406 / 11,278 |
| 80-97.5 | 1.90 (1.73-2.08) | $4.44 \times 10^{-43}$ | 719 / 2,981 |
| >97.5 | 2.99 (2.52-3.54) | $1.44 \times 10^{-36}$ | 151 / 378 |
| **AF PRS** | | | |
| <2.5 | 0.44 (0.26-0.73) | 0.002 | 15 / 478 |
| 2.5-20 | 0.60 (0.50-0.72) | $1.69 \times 10^{-8}$ | 143 / 3,303 |
| 20-80 | 1 (reference) | - | 779 / 11,035 |
| 80-97.5 | 1.94 (1.72-2.19) | $4.28 \times 10^{-27}$ | 406 / 3,039 |
| >97.5 | 3.19 (2.56-3.98) | $8.67 \times 10^{-25}$ | 88 / 405 |
| **Breast cancer PRS** | | | |
| <2.5 | 0.43 (0.16-1.16) | 0.09 | 4 / 280 |
| 2.5-20 | 0.44 (0.30-0.65) | $3.58 \times 10^{-5}$ | 29 / 1,954 |
| 20-80 | 1 (reference) | - | 221 / 6,577 |
| 80-97.5 | 1.94 (1.56-2.42) | $3.90 \times 10^{-9}$ | 123 / 1,860 |
| >97.5 | 3.05 (2.04-4.55) | $5.34 \times 10^{-8}$ | 27 / 257 |
| **Prostate cancer PRS** | | | |
| <2.5 | 0.10 (0.01-0.73) | 0.02 | 1 / 256 |
| 2.5-20 | 0.38 (0.25-0.58) | $4.90 \times 10^{-6}$ | 25 / 1,770 |
| 20-80 | 1 (reference) | - | 233 / 5,921 |
| 80-97.5 | 2.14 (1.74-2.64) | $6.58 \times 10^{-13}$ | 146 / 1,649 |
| >97.5 | 3.93 (2.79-5.53) | $3.90 \times 10^{-15}$ | 39 / 218 |

CHD = coronary heart disease, AF = atrial fibrillation or flutter, T2D = type 2 diabetes, PRS = polygenic risk score
